# High-quality SNPs from genic regions highlight introgression patterns among European white oaks (*Quercus petraea* and *Q. robur*)

**DOI:** 10.1101/388447

**Authors:** Tiange Lang, Pierre Abadie, Valérie Léger, Thibaut Decourcelle, Jean-Marc Frigerio, Christian Burban, Catherine Bodénès, Erwan Guichoux, Grégoire Le Provost, Cécile Robin, Naoki Tani, Patrick Léger, Camille Lepoittevin, Veronica A. El Mujtar, François Hubert, Josquin Tibbits, Jorge Paiva, Alain Franc, Frédéric Raspail, Stéphanie Mariette, Marie-Pierre Reviron, Christophe Plomion, Antoine Kremer, Marie-Laure Desprez-Loustau, Pauline Garnier-Géré

## Abstract

In the post-genomics era, non-model species like most *Fagaceae* still lack operational diversity resources for population genomics studies. Sequence data were produced from over 800 gene fragments covering ~530 kb across the genic partition of European oaks, in a discovery panel of 25 individuals from western and central Europe (11 *Quercus petraea*, 13 *Q. robur*, one *Q. ilex* as an outgroup). Regions targeted represented broad functional categories potentially involved in species ecological preferences, and a random set of genes. Using a high-quality dedicated pipeline, we provide a detailed characterization of these genic regions, which included over 14500 polymorphisms, with ~12500 SNPs −218 being triallelic-, over 1500 insertion-deletions, and ~200 novel di- and tri-nucleotide SSR loci. This catalog also provides various summary statistics within and among species, gene ontology information, and standard formats to assist loci choice for genotyping projects. The distribution of nucleotide diversity (*θπ*) and differentiation (*F*_*ST*_) across genic regions are also described for the first time in those species, with a mean n *θπ* close to ~0.0049 in *Q. petraea* and to ~0.0045 in *Q. robur* across random regions, and a mean *F*_*ST*_ ~0.13 across SNPs. The magnitude of diversity across genes is within the range estimated for long-term perennial outcrossers, and can be considered relatively high in the plant kingdom, with an estimate across the genome of 41 to 51 million SNPs expected in both species. Individuals with typical species morphology were more easily assigned to their corresponding genetic cluster for *Q. robur* than for *Q. petraea*, revealing higher or more recent introgression in *Q. petraea* and a stronger species integration in *Q. robur* in this particular discovery panel. We also observed robust patterns of a slightly but significantly higher diversity in *Q. petraea*, across a random gene set and in the abiotic stress functional category, and a heterogeneous landscape of both diversity and differentiation. To explain these patterns, we discuss an alternative and non-exclusive hypothesis of stronger selective constraints in *Q. robur*, the most pioneering species in oak forest stand dynamics, additionally to the recognized and documented introgression history in both species despite their strong reproductive barriers. The quality of the data provided here and their representativity in terms of species genomic diversity make them useful for possible applications in medium-scale landscape and molecular ecology projects. Moreover, they can serve as reference resources for validation purposes in larger-scale resequencing projects. This type of project is preferentially recommended in oaks in contrast to SNP array development, given the large nucleotide variation and the low levels of linkage disequilibrium revealed.

## Introduction

High-throughput techniques of the next-generation sequencing (NGS) era and increased genome sequencing efforts in the last decade have greatly improved access to genomic resources in non-model forest tree species (Neale and Kremer 2011, Neale *et al*. 2013; Plomion *et al*. 2016). However, these have only been applied recently to large-scale ecological and population genomics research (Holliday *et al*. 2017). One notable exception are studies undertaken in the model genus *Populus* (e.g. Zhou *et al*. 2014, Geraldes *et al*. 2014, Christe *et al*. 2016b) that benefited from the first genome sequence completed in 2006 in *P. trichocarpa* (Tuskan *et al*. 2006). In *Fagaceae*, previous comparative mapping and “omics” technologies (reviewed in Kremer *et al*. 2012) with recent development of genomic resources (e.g. Faivre-Rampant *et al*. 2011; Tarkka *et al.* 2013; Lesur *et al*. 2015; Lepoittevin *et al*. 2015, Bodénès *et al*. 2016) set the path to very recent release of genome sequences to the research community (*Quercus lobata,* Sork *et al.* 2016; *Q. robur,* Plomion *et al*. 2016, 2018; *Q. suber,* Ramos *et al*. 2018; *Fagus sylvatica*, Mishra *et al*. 2018), and these provide great prospects for future evolutionary genomics studies (Petit *et al*. 2013; Parent *et al*. 2015; Cannon *et al*. 2018; Lesur *et al*. 2018).

Recently, building from the European oaks genomic resources (*Quercus Portal* at https://arachne.pierroton.inra.fr/QuercusPortal/ and references therein), natural populations of 4 *Quercus* species (*Q. robur, Q. petraea, Q. pyrenaica, Q. pubescens*) were genotyped for ~4000 single-nucleotide polymorphisms (SNPs, from an initial 8K infinium array, Lepoittevin *et al*. 2015). The data were further analysed (Leroy *et al*. 2017), with results extending previous knowledge on their likely diversification during glacial periods, as well as their recolonization history across Europe and recent secondary contacts (SC) after the last glacial maximum (Hewitt 2000; Petit *et al*. 2002a; Brewer *et al*. 2002). Using recent model-based inference allowing for heterogeneity of migration rates (Roux *et al*. 2014; Tine *et al.* 2014), Leroy *et al*. (2017) showed that the most strongly supported demographic scenarios of species diversification, allowing for gene flow among any pair of the four species mentioned above, included very recent SC, due to a much better fit for patterns of large heterogeneity of differentiation observed across SNP loci (confirmed by Leroy *et al*. 2019, using ~15 times more loci across the genome and the same inference strategy). These recent SC events have been documented in the last decade in many patchily distributed hybrid zones where current *in situ* hybridization can occur among European oak species (e.g. Curtu *et al*. 2007; Jensen *et al.* 2009; Lepais and Gerber 2011; Guichoux *et al*. 2013). The resulting low levels of differentiation among *Q. robur* and *Q. petraea* in particular is traditionally linked to a model of contrasted colonization dynamics, where the second-in-succession species (*Q. petraea*) is colonizing populations already occupied by the earlier pioneering *Q. robur* (Petit *et al.* 2003). This model predicts asymmetric introgression towards *Q. petraea* (see Currat *et al*. 2008), as often observed in interspecific gene exchanges (Abbott *et al*. 2003), and a greater diversity in *Q. petraea* was documented at SNP loci showing higher differentiation (Guichoux *et al.* 2013). The directionality of introgression in oaks was also shown to depend on species relative abundance during mating periods in particular stands (Lepais *et al*. 2009, 2011). Nethertheless, oaks like other hybridizing taxa are known for the integration of their species parental gene pools and strong reproductive isolation barriers (Muir *et al*. 2000; Muir and Schlötterer 2005; Abadie *et al.* 2012, Lepais *et al*. 2013; Ortiz-Barrientos and Baack 2014; Christe *et al*. 2016a), raising essential questions about the interacting roles of divergent (or other types of) selection, gene flow, and recombination rates variation in natural populations, and their imprints on genomic molecular patterns of variation (e.g. Zhang *et al*. 2016; Christe *et al*. 2016b; Payseur and Rieseberg 2016).

These issues will be better addressed with genome-wide sequence data in many samples (Buerkle *et al*. 2011), which will be facilitated in oaks by integrating the newly available genome sequence of *Quercus robur* to chosen HT resequencing methods (Jones and Good 2016; e.g. Zhou and Holliday 2012; Lesur *et al.* 2018 for the first target sequence capture study in oaks). However, obtaining high quality haplotype-based data required for nucleotide diversity estimation and more powerful population genetics inferences will likely require the development of complex bioinformatics pipelines dedicated to high heterozygosity genomes and solid validation methods for polymorphism detection (e.g. Geraldes *et al*. 2011; Christe *et al*. 2016b).

Therefore, the objectives of this work were first to provide a detailed characterization of sequence variation in *Quercus petraea* and *Quercus robur*. To that end, we validated previous unpublished sequence data from the classical Sanger’ chain-terminating dideoxynucleotides method (Sanger *et al*. 1977). These sequences targeted fragments of gene regions in a panel of individuals sampled across the western and central European part of both species geographic range. Both functional and expressional candidate genes potentially involved in species ecological preferences, phenology and host-pathogen interactions were targeted, as well as a reference set of fragments randomly chosen across the last oak unigene (Lesur *et al*. 2015). These data were obtained within the framework of the EVOLTREE network activities (http://www.evoltree.eu/). Second, we aimed at estimating the distributions of differentiation and nucleotide diversity across these targeted gene regions for the first time in those species, and further test the robustness of comparative diversity patterns observed in the context of both species contrasted dynamics and introgression asymmetry. We discuss the quality, representativity and usefulness of the resources provided for medium scale genotyping landscape ecology projects or as a reference resource for validation purposes in larger-scale resequencing projects.

## Material and Methods

### Sample collection

The discovery panel (*DiP*) included 25 individuals from 11 widespread forest stands with 2 to 4 individuals per location (13 from *Q. robur,* 11 from *Q. petraea*, 1 from *Q. ilex* to serve as outgroup, in Table 1).

**Table 1.**
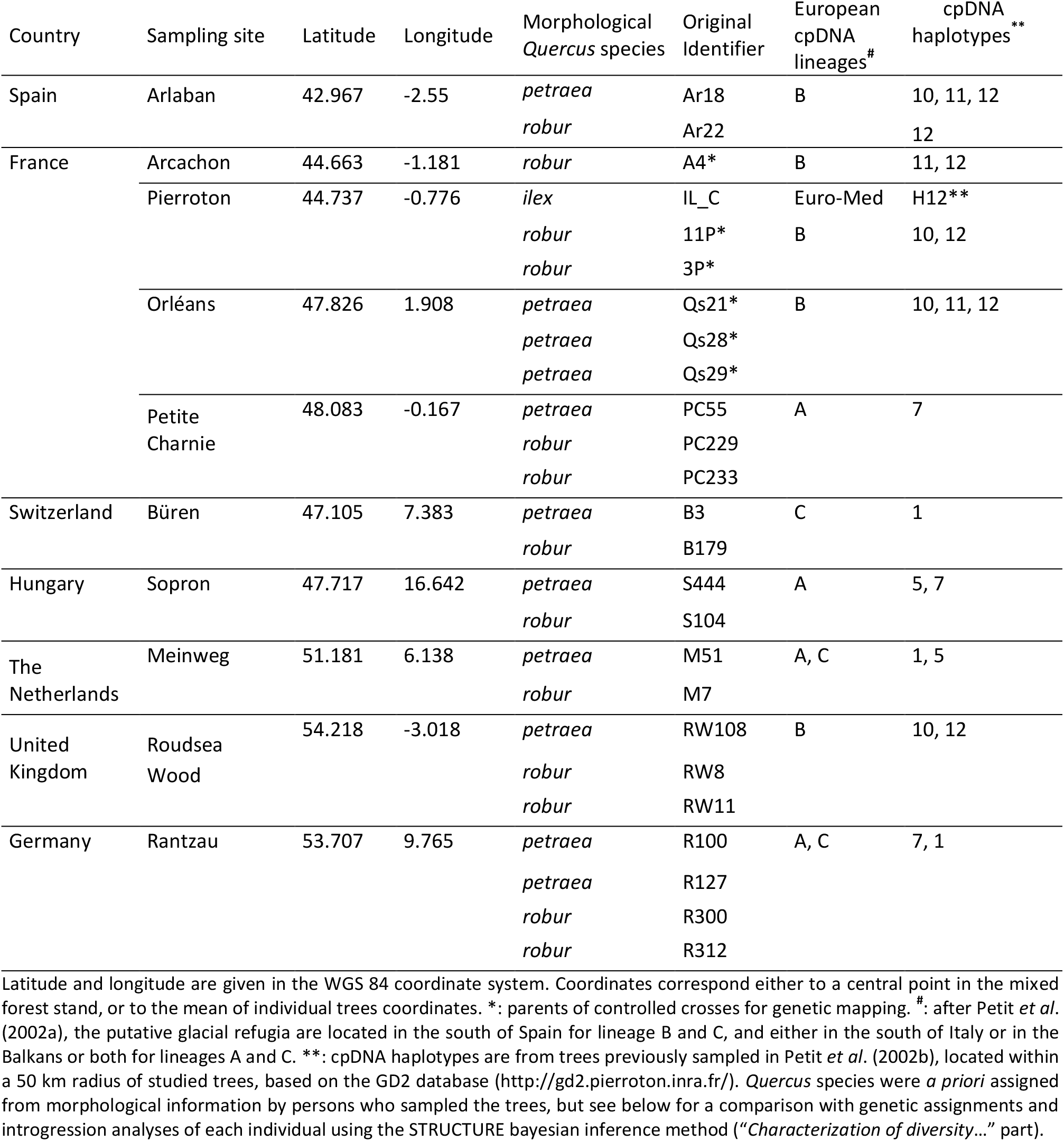
Geographic location of 25 sampled individuals from *Quercus petraea*, *Q. robur* and *Q. ilex.*

| Country | Sampling site | Latitude | Longitude | Morphological<br>*Quercus* species | Original<br>Identifier | European<br>cpDNA<br>lineages<sup>#</sup> | cpDNA<br>haplotypes<sup>**</sup> |
| --- | --- | --- | --- | --- | --- | --- | --- |
| Spain | Arlaban | 42.967 | -2.55 | *petraea* | Ar18 | B | 10, 11, 12 |
|  |  |  |  | *robur* | Ar22 |  | 12 |
| France | Arcachon | 44.663 | -1.181 | *robur* | A4* | B | 11, 12 |
|  | Pierroton | 44.737 | -0.776 | *ilex* | IL_C | Euro-Med | H12** |
|  |  |  |  | *robur* | 11P* | B | 10, 12 |
|  |  |  |  | *robur* | 3P* |  |  |
|  | Orléans | 47.826 | 1.908 | *petraea* | Qs21* | B | 10, 11, 12 |
|  |  |  |  | *petraea* | Qs28* |  |  |
|  |  |  |  | *petraea* | Qs29* |  |  |
|  | Petite<br>Charnie | 48.083 | -0.167 | *petraea* | PC55 | A | 7 |
|  |  |  |  | *robur* | PC229 |  |  |
|  |  |  |  | *robur* | PC233 |  |  |
| Switzerland | Büren | 47.105 | 7.383 | *petraea* | B3 | C | 1 |
|  |  |  |  | *robur* | B179 |  |  |
| Hungary | Sopron | 47.717 | 16.642 | *petraea* | S444 | A | 5, 7 |
|  |  |  |  | *robur* | S104 |  |  |
| The<br>Netherlands | Meinweg | 51.181 | 6.138 | *petraea* | M51 | A, C | 1, 5 |
|  |  |  |  | *robur* | M7 |  |  |
| United<br>Kingdom | Roudsea<br>Wood | 54.218 | -3.018 | *petraea* | RW108 | B | 10, 12 |
|  |  |  |  | *robur* | RW8 |  |  |
|  |  |  |  | *robur* | RW11 |  |  |
| Germany | Rantzau | 53.707 | 9.765 | *petraea* | R100 | A, C | 7, 1 |
|  |  |  |  | *petraea* | R127 |  |  |
|  |  |  |  | *robur* | R300 |  |  |
|  |  |  |  | *robur* | R312 |  |  |

Latitude and longitude are given in the WGS 84 coordinate system. Coordinates correspond either to a central point in the mixed forest stand, or to the mean of individual trees coordinates. \*: parents of controlled crosses for genetic mapping. <sup>#</sup>: after Petit *et al.* (2002a), the putative glacial refugia are located in the south of Spain for lineage B and C, and either in the south of Italy or in the Balkans or both for lineages A and C. \*\*: cpDNA haplotypes are from trees previously sampled in Petit *et al.* (2002b), located within a 50 km radius of studied trees, based on the GD2 database (http://gd2.pierroton.inra.fr/). *Quercus* species were *a priori* assigned from morphological information by persons who sampled the trees, but see below for a comparison with genetic assignments and introgression analyses of each individual using the STRUCTURE bayesian inference method (“*Characterization of diversity...*” part).

These stands occur across a large part of both *Quercus* species natural distributions, spanning ~20° in longitude (~2200 km) and ~11° in latitude (~1250 km) in western and central Europe (Fig. S1, Supporting Information). They are also located in areas covering the three major cpDNA lineages A, B and C (among five) that indicate different historical glacial refugia (Petit *et al.* 2002a), and extend much further geographically towards northern, eastern and south-eastern European borders (Table 1, after Petit *et al*. 2002b). One stand (Sopron, in Hungary), also occurs within the large geographic distribution of the most Eastern lineage E, in a region where lineages A and C also occur. Individuals were chosen either on the basis of their differing leaf morphology among *Q. robur* and *Q. petraea* species (Kremer *et al*. 2002a), or as parents of mapping pedigrees (e.g. Bodénès *et al*. 2016, see Table 1).

Leaves were sampled, stored in silica gel and sent to INRA (Cestas, France) for DNA extraction following Guichoux *et al*. (2013). DNA quality and concentration were assessed with a Nanodrop spectrophotometer (NanoDrop Technologies, Wilmington, 152 DE, USA) and by separating samples in 1% agarose gels stained with ethidium bromide. Extractions were repeated until we obtained at least 20 micrograms of genomic DNA per sample, which was needed for a few thousands individual PCRs.

### Choice of genic regions for resequencing

Genic regions were chosen from over 103 000 Sanger sequences available in expressed sequence tags (EST) databases at the start of the project. These sequences corresponded to 14 cDNA libraries that were prepared with many individuals from both species. They were assembled before finally selecting 2000 fragments for resequencing (Appendix S1 and Fig. S2-A, Supporting information for more on methods producing the original working assembly (*orict*); see also Ueno *et al*. 2010). The targeted fragments were chosen from an extensive compilation of both expressional and functional candidate genes that would likely be involved in white oaks’ divergent functions and/or local adaptation, using model and non-model species databases or published results (see Appendix S1 and Fig. S2-B, Supporting information for more details on the strategy followed, and Table S1 for designed primers).

### Data production and polymorphism discovery in resequenced fragments

All the sequencing work was performed by Beckman Coulter (Agencourt Bioscience Corporation, Beverly, MA, USA) on ABI3730 capillary sequencers (Applied Biosciences) after preparing DNA samples according to the company’s guidelines. Various data quality steps were followed for maximizing the amount and quality of the sequences finally obtained (Fig. 1-A, and Appendix S1, Supporting information for further analyses across 2000 amplicons).

**Figure 1.**
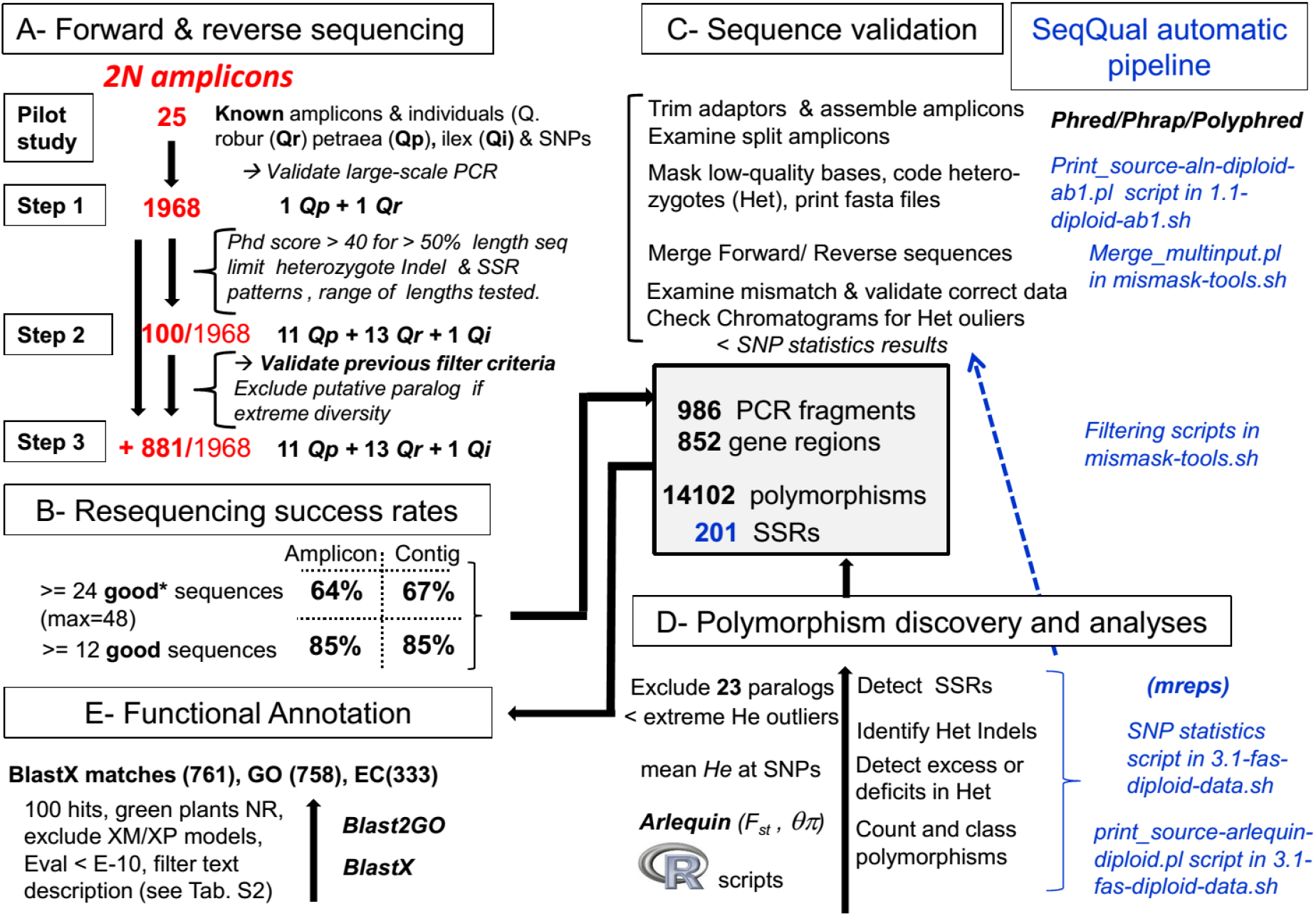
Bioinformatics strategy for sequence data production, amplicon assembly, functional annotation, and polymorphism discovery. Scripts used are in italics (see text for further details). GO: Gene Ontology, EC: Enzyme Commission ID. * A **good** sequence is defined as having a minimum of 50% of its nucleotides with a Phred score above 30.

Text Forward and reverse sequences were produced for 986 amplicons across 25 individuals (100+881 in steps 2 and 3, Fig. 1-A), and more than 85% of them yielded at least 12 high-quality sequences (Fig. 1-B and column L in Table S1, Supporting information). All amplicon assembly steps, merging, trimming, and filtering/masking based on quality were performed with our *SeqQual* pipeline (https://github.com/garniergere/SeqQual), with examples of data and command files. This repository compiles and extends former work dealing with 454 data (Brousseau *et al*. 2014; El Mujtar *et al.*2014), providing Bioperl scripts used here that automatically deal with Sanger haploid or diploid DNA sequences and allow fasta files post-processing in batch (Fig. 1-C). Sequence variants discovery was finally performed using an error rate below 0.001 (i.e. Phred score above 30, Ewing *et al*. 1998, and see Appendix S1, Supporting information for more details).

Simple sequence repeat (SSR) patterns were further detected or confirmed from consensus sequences using the *mreps* software (Kolpakov *et al.* 2003; see Fig. 1-D, and https://github.com/garniergere/Reference.Db.SNPs.Quercus/ for a R script parsing *mreps* output). Various additional steps involving the treatment of insertion-deletion polymorphisms (indels) and heterozygote indels (*HI*) in particular, allowed missing data from polymorphic diploid sequence to be minimized (see Appendix S1, Supporting information).

### Functional annotation

Resequenced genic regions were annotated using the BlastN best hits of their corresponding *orict* contigs and those of their expected amplicons (*orict-cut*) to most recent oak assembly (*ocv4*, Lesur *et al*. (2015); see Table S2-C, Supporting information). Final consensus sequences for these regions originated from both *orict* and *ocv4* (396 and 368 respectively, see Table S2-A, S2-B, and Appendices S1 and S3, Supporting information), aiming at retrieving the longest sequences, while avoiding to target those with possible chimeric sequences. Functional annotation was then performed via homology transfer using BlastX 2.6.0+program at NCBI (https://blast.ncbi.nlm.nih.gov/Blast.cgi) with parameters to optimize speed, hits’ annotation description and GO content (Fig. 1-E and Table-S2, Supporting information). Retrieval of GO terms were performed with Blast2GO (Conesa *et al.* 2005 free version at https://www.blast2go.com/blast2go-pro/b2g-register-basic) and validation of targeted annotations with Fisher Exact enrichment tests (details of Blast2GO analyses provided in Appendix S1, Supporting information).

### Characterization of diversity and genetic clustering

Using the *SNP-stats* script for diploid data from *SeqQual*, simple statistics were computed across different types of polymorphisms (SNPs, indels, SSRs…) including minimum allele frequencies (*maf*) and heterozygote counts, Chi-square tests probability for Hardy-Weinberg proportions, *G*_*ST*_ (Nei 1987) and *G_ST_’* standardized measure (Hedrick 2005). Complex polymorphisms (involving heterozygote indels (*HI*) and/or SSRs,) were also further characterized (see Appendix S1, Supporting information), and data formatted or analyzed using either Arlequin 3.5 (Excoffier and Lischer 2010), *SeqQual* (e.g. for Arlequin input file with phase unknown, Fig. 1-C), or R scripts. Nucleotide diversity *θπ* (Nei 1987), based on the average number of pairwise differences between sequences, and its evolutionary variance according to Tajima (1993), were also estimated and compared among species and across candidate genes grouped by broad functional categories (see column F in Table S1, Supporting information), and Weir and Cockerham (1984) *F*_*ST*_ estimates of differentiation were computed among species for SNP data along genic regions using analyses of molecular variance (Excoffier 2007).

The initial morphological species samples were compared to the genetic clusters obtained with the Structure v2.3.3 inference method (Falush *et al.* 2003) in order to test possible levels of introgression across individuals. We used the admixture model allowing for mixed ancestry and the correlated allele frequencies assumption for closely related populations as recommended defaults, and since they best represent previous knowledge on each species genetic divergence across their range (e.g. Guichoux *et al.* 2013). Preliminary replicate runs using the same sample of loci produced very low standard deviation across replicates of the data log likelihood given K (ln Pr(X/K), see Fig. S3-A, Supporting information). We thus resampled loci at random for each of 10 replicate datasets in 3 different manners to add genetic stochasticity: 1) one per region, 2) one per 100 bp block, and 3) one per 200 bp block along genes (see Appendix S1, Supporting information and https://github.com/garniergere/Reference.Db.SNPs.Quercus/tree/master/STRUCTURE.files for examples of Structure files as recommended by Gilbert *et al.* (2012), along with R scripts for outputs). Statistical independence among loci within each species was verified with Fisher’s exact tests implemented in Genepop 4.4 (Rousset 2008).

## Results

### Polymorphisms typology and counts

Among the amplicons tested, 986 were successful, 13 did not produce any data and 23 were excluded because of paralog amplifications (Fig. 1-C and Table S1, Supporting information). Around 25% of the successful amplicons overlapped and were merged, consistently with their original design across contigs. Despite the presence of *HI* patterns due to SSR or indels, most amplicons were entirely recovered with forward and reverse sequencing. Several were however kept separate (5% of the total), either because of functional annotation inconsistency, or because amplicon overlap was prevented by the presence of SSRs or putative large introns (see “Final gene region ID” column with −F/−R suffix in Table S1, Supporting information). We finally obtained 852 genic regions covering in total ~529 kilobases (kb), with an average size of 621 bp per region, ranging from 81 to 2009 bp (Table 2, and Appendix S4, Supporting information, for genomic consensus sequences).

**Table 2.** STypology of polymorphisms in successfully resequenced amplicons.

|  | Both species<br>and<br>introgressed<br>individuals | *Q. petraea* | *Q. robur* | *Q. ilex* |
| --- | --- | --- | --- | --- |
| Total length resequenced (bp) | 529281 | - | - | 196676 |
| Number (Nb) of amplicons | 986 | - | - | 486 |
| Nb of genic regions | 852 | - | - | 394 |
| Mean genic region size - N50 size (bp) | 621-700 | - | - | 500-539 |
| Minimum - Maximum genic region size (bp) | 81-2009 | - | - | 198-1285 |
| Estimated intron sequences (bp) | 186827 | - | - | - |
| Mean haploid sample size (total sequence) | 34.71 | 13.35 | 18.28 | - |
| **Polymorphism in 852 genic regions** |  |  |  |  |
| Mean haploid sample size (variants) | 32.16 | 12.57 | 13.85 | - |
| Monomorphic genic regions | 15 (1.76%) | 18 (2.14%) | 21 (2.52%) | - |
| Genes with at least one single base indel | 591 | 345 | 379 | - |
| " " " " one larger indel (>1 bp) | 252 | 190 | 214 | - |
| " " " " one SSR (>=di) | 163 | - | - | - |
| SNPs only (excluding 1 bp indels) | 12478 | 7511 | 8078 | - |
| Indels (1 bp) | 1213 | 751 | 809 | - |
| Indels (2-5 bp) | 221 | 142 | 161 | - |
| Indels ( 6-10 bp) | 88 | 72 | 71 | - |
| Indels ( 11-50 bp, excl. SSRs) | 98 | 81 | 79 | - |
| Indels (74,146,219,341 bp, excl. SSRs) | 4 | 3 | 4 | - |
| **Total number of polymorphisms** | **14102** | **8560** | **9202** | **676** |
| *Triallelic SNPs* | *218* | *141* | *165* | - |
| *...Singletons (incl. 1 bp indels)* | *4334* | *1990* | *2151* | - |
| *...Variable SSRs (excl. homopolymers)* | *111* | - | - | - |
| Total length with sequence variant positions | 17594 | 10765 | 11451 | - |
| Sequence length of indels and complex polymorphisms (Indels and SSRs) | 5116 | - | - | - |

Compared to the EST-based expected total fragment size of ~ 357 kb, around 187 kb of intron sequence was recovered across 460 of the resequenced regions (assuming intron presence if an amplicon size was above its expected size by 40 bp). Introns represented ~35% of genic regions in length and ~51% of those including introns.

We observed 14102 polymorphisms in both species across 852 gene regions, 15 of those regions (<2%) being monomorphic (Table 2). This corresponds to 1 polymorphism per ~38 bp, or 1 per ~30 bp when considering the total number of variant positions in both species (17594 bp, Table 2). Remarkably, variant positions involving larger indels, SSRs and mixed complex polymorphism patterns represented ~30% of the total variant positions (Table 2, and see their exhaustive lists with various statistics in Table S3 and S4, Supporting information). We observed 12478 SNPs (88.5% of all polymorphisms), 1 SNP per 42 bp, and 218 triallelic SNPs (~1.75% of SNPs) were confirmed by visual examination of chromatograms.

Considering only one species, we observed on average 1 variant position per ~48 bp, 1 polymorphism per ~60 bp, and 1 SNP per ~68 bp. Among indels, 1213 (8.6% of all polymorphisms) were single base, 309 ranged from 2 to 10 bp, and 102 had sizes above 10 bp which were mostly shared among species (Table 2). In this range-wide sample, there were 4334 singletons among all single base polymorphisms, 506 of them being indels. Overall, indels were present in 69% of gene regions and non-single base ones across ~30% of them. Excluding homopolymers (see Appendix S1, Supporting information), we detected 201 SSRs occurring on 163 gene regions by considering a minimum repeat numbers of 4 and a mismatch rate among repeats below 10% (Table 2, Table S1 and Table S5, Supporting information), and 55% (111) were polymorphic in our sample of individuals (Table 2). Among them, 89 (44%) had dinucleotide repeats and 65 (32%) trinucleotide repeats. The SSRs with the lowest number of repeats (<5) had a majority (59%) of repeat sizes between 4 and 7, the rest being trinucleotides (Table S5, Supporting information).

Using the same PCR conditions, homologous sequence data were obtained for one individual of the outgroup *Quercus ilex* across 37% of the gene regions (~197 kb, 397 sequences, 676 heterozygous sites in Table 2), which illustrates both their sequence similarity yet divergence for a species belonging to the *Ilex* versus *Quercus* taxonomic group (Lepoittevin *et al*. 2015; see Table S1 column Q, and see Appendix S5, Supporting information, for *Q. ilex* genomic sequences).

### Annotations and GO term distributions

BlastX matches with *E*-values below 10^−30^ were found for ~97% (738/764) of the contig consensus, only 11 sequences (1.4%) having hits with *E*-values above 10^−10^ that were all among the reference random sample (see BlastX criteria in Table S2, Supporting information). The most represented species among the best hits with informative annotations were *Prunus persica (111)*, *Theobroma cacao (91)*, *Morus notabilis (57)* and *Populus trichocarpa (45)* (Appendix S6-A, Supporting information), which probably illustrates both the close phylogenetic relationships among *Quercus* and *Prunus* genera, consistently with results obtained on the larger *ocv4* assembly (Lesur *et al.* 2015), and the quality and availability of *P. persica* genome annotation (Verde *et al*. 2013, 2017).

Between 1 to 30 GO terms could be assigned to 761 sequences, with EC codes and InterProScan identifiers for 343 and 733 of them respectively (Fig. 1, and Table S2, Supporting information). The most relevant GO terms were then retained using the Blast2GO “annotation rule” (Conesa *et al*. 2005) that applies filters from the Direct Acyclic Graph (DAG) at different levels (Fig. 2, Fig. S4-A- to-F, Supporting information).

**Figure 2.**
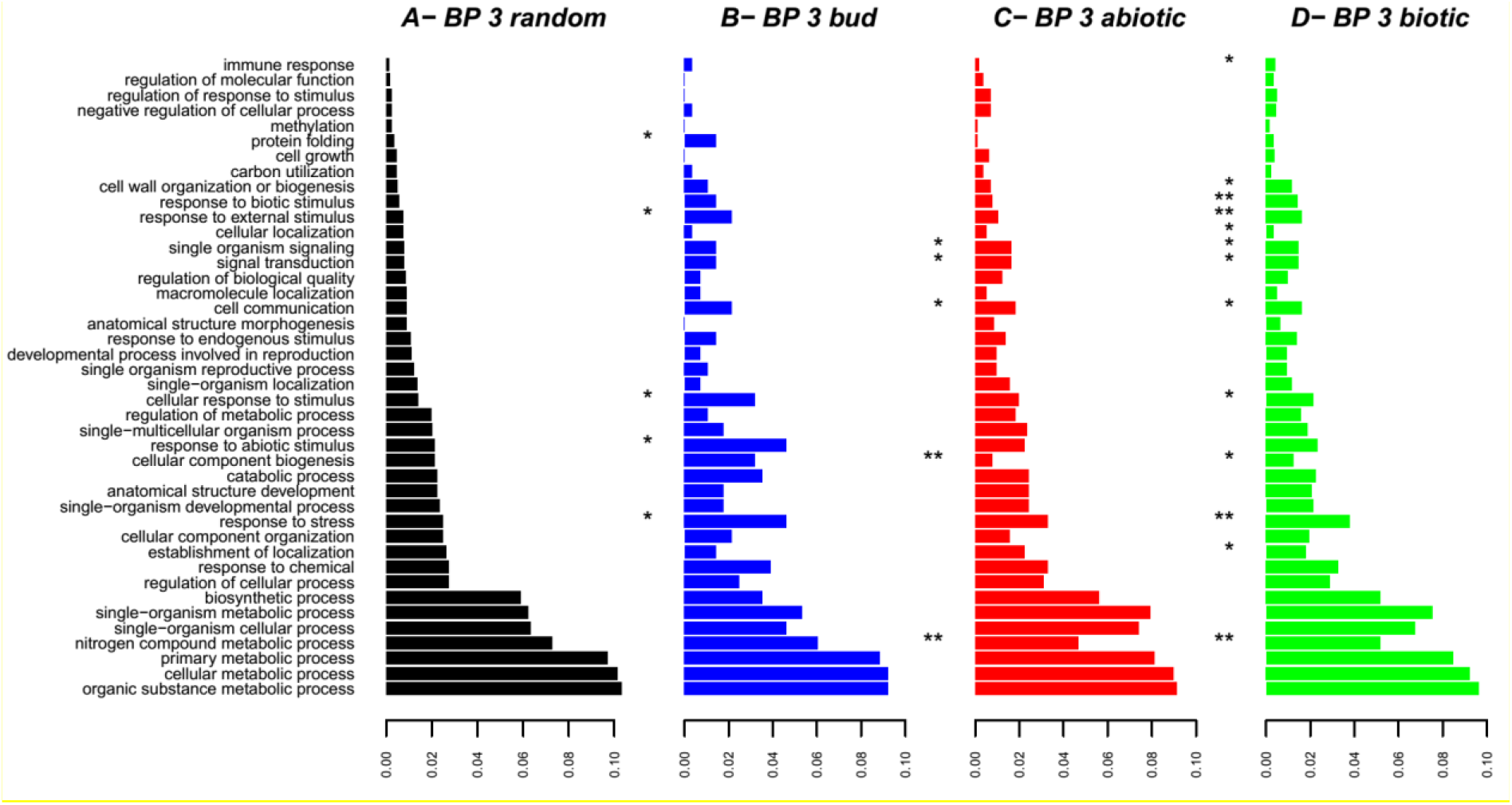
Distributions of GO terms across different gene lists (*bud*, *abiotic* and *biotic*) at biological process level 3, and Fisher exact tests across pairs of sequence clusters with the same GO terms between the random list and other lists. Significance levels *: P<0.05, **: P<0.01.

At biological process (BP) level 3, apart from general terms involving “metabolic processes”, a large number of sequences (between ~100 and ~150) were mapped to “response to…” either “…stress”, “…abiotic stimulus” or “…chemical”, and also to categories linked to developmental processes (Fig. S4-D, Supporting information).

Enrichment tests also revealed a significant increase at both BP levels 2 and 3 for the following GO categories: “response to stress” or “external stimulus” for *bud* and *biotic* gene lists, “response to abiotic stimulus” for the *bud* list, and “immune” and “biotic stimulus” responses for the *biotic* list (see Fig. 2-B to 2-D compared to Fig. 2-A, and Fig. S5, Supporting information). Most of these exact tests (>80%) were still significant when selecting genes attributed exclusively to one particular list (in Table S1, Supporting information), which adds to the relevance of our original gene lists in targeting particular functional categories.

### Species assignment and introgressed individuals

In both species, the proportion of significant association tests among the loci used for clustering (> two million within each species) was generally one order of magnitude below the type-I error rates at 5% or 1%. This indicates a very low background LD within species at their range levels, consistently with the underlying model assumptions used in Structure. Based on both ln Pr(X/K) ⍰K statistics and as expected, the optimal number of genetic clusters inferred was 2, whatever the number of polymorphisms and type of sampling (Fig. 3, Fig. S3 and S6, Supporting information).

**Figure 3.**
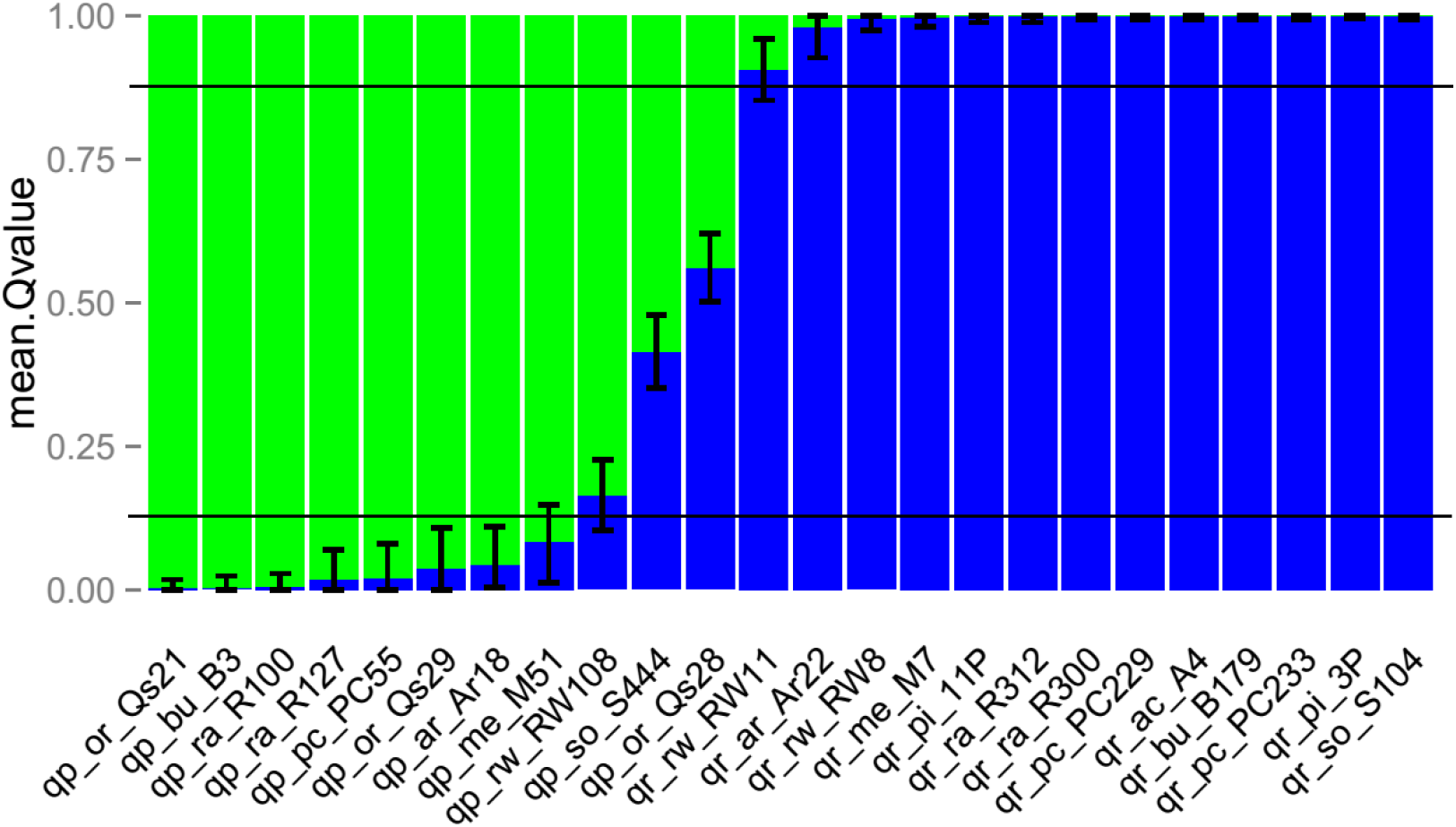
Posterior assignment probabilities of individuals into two optimal clusters from STRUCTURE analyses, sorted in increasing order of belonging to cluster 2 (*Q. robur* (qr-), in blue here, the alternative cluster 1 matching *Q. petraea* (qp-) in green), apart from individuals with higher introgression levels. Each bar represents one individual and includes mean upper and lower bounds of 90% Bayesian confidence intervals around mean *Q*-values across 10 replicates. Each replicate is a different random sample of 1785 polymorphisms. Horizontal black lines represent the 0.125 and 0.875 values, which can be considered as typical thresholds for back-crosses and later-generation hybrids (Guichoux *et al*. 2013), values within those thresholds suggesting a mixed ancestry with the other species for a small number of generations in the past.

Most individuals (20) clearly belonged to either cluster with a mean probability of cluster assignment above 0.9, which was not significantly different from 1, based on mean values of 90% Bayesian credible intervals (BCI) bounds across replicates, and for different types of sampling or SNP numbers (Fig. 3 and Fig. S6, Supporting information). Two individuals from Roudsea Wood in UK, the most northerly forest stand of this study, were considered to be significantly introgressed, each from a different cluster, since both showed a BCI that did not include the value “1” across other replicated runs and SNP sampling (Fig. S6, Supporting information), RW108 also having a mean probability above 0.125 (Fig. 3). Although M51 has a mean assignment value close to that of RW11 in the particular run shown in Fig.3, its BCI was larger and often included the zero value in other runs (Fig. S6, Supporting information), so it was assigned to the *Q. petraea* cluster. In the initial morphological *Q. petraea* group, two individuals were also clearly of recent mixed ancestry: one from the easternmost forest stand of Sopron (S444), and another one (Qs28) from central France, considered previously to be a *Q. petraea* parental genotype in two oak mapping pedigrees (Bodénès *et al*. 2012, 2016; Lepoittevin *et al.* 2015). However, Qs28 shows here a clear F1 hybrid pattern, given its probability values close to 0.5 and its BCI maximum upper and minimum lower bound values of 0.30 and 0.61 respectively across runs (Fig. 3 and Fig. S6-A to S6-J, Supporting information). Testing 3 or 4 possible clusters showed the same ancestry patterns for the introgressed individuals with 2 main clusters and similar *Q*-values (data not shown), which does not support alternative hypotheses of introgression from different species in those individuals.

### Large heterogeneity of diversity and differentiation across genes

Nucleotide diversity was thus estimated in each parental species after excluding Qs28, RW108, S444 and RW11, which were considered to be the 4 most introgressed individuals (see Fig. 3 above). We then checked how the remaining samples represented species’ diversity. Starting with one individual, we observe a dramatic drop in the mean proportion of new variant positions brought by each new individual in any species (*Mpn*) as a function of the initial sample size, followed by a subsequent stabilization (Fig. 4-A, and see Fig. S7-A, Supporting information). Indeed, *Mpn* was only around 11% when going from 4 to 5 individuals in both species, and stabilized below 5% after 8 individuals in *Q. robur* (Fig. 4-A). We thus decided to retain 726 gene regions with at least 8 gametes per species (listed in column L in Table S1, Supporting information). The larger *Q. robur* sample after excluding the most introgressed individuals (24 versus 16 gametes in *Q. petraea*) only exhibited slightly higher polymorphism counts than in *Q. petraea* overall (Table 3).

**Figure 4.**
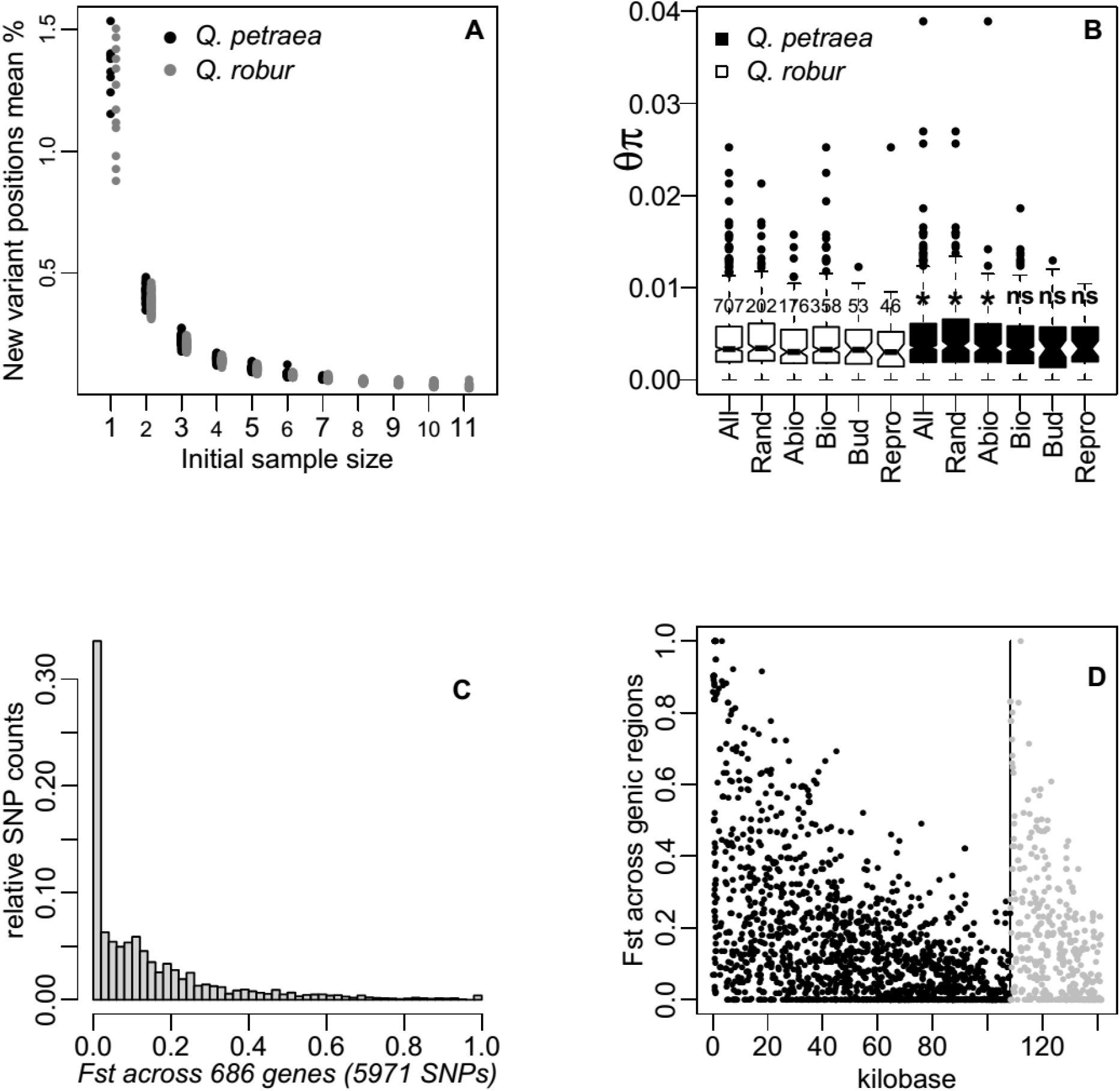
Mean proportion of new variant sites brought by each new distinct individual added to all possible initial sample size combinations (A); Mean nucleotide diversity (considering all polymorphisms) in both species across genic regions, and different functional categories (B) compared between species with Wilcoxon signed-rank tests: significant at Pr<5% (*), non-significant (ns); Histogram of *Fst* estimates across polymorphic gene regions with a minimum of 8 gametes per species, after excluding singletons and grouping negative with null values (C); Manhattan plot of *Fst* estimates sorted by mean *Fst* values across randomly chosen (black dots) and Bud phenology (grey dots) genic regions (D).

**Table 3.** Polymorphism counts and nucleotide diversity in parental species across genic regions with larger sample sizes.

| Polymorphism in 726 gene fragments | both species | *Q. petraea* | *Q. robur* |
| --- | --- | --- | --- |
| Number of individuals considered | 20 | 8 | 12 |
| Monomorphic gene fragments | 17 (2.34%) | 19 (2.63%) | 20 (2.87%) |
| Total number of polymorphisms | 11089 | 7061 | 7721 |
| SNPs only | 9867 | 6226 | 6830 |
| All Indels and SSRs | 1222 | 835 | 891 |
| Exclusive polymorphisms | - | 3359 | 4024 |
| Singletons among them (%) | - | 0.456 | 0.437 |
| Shared polymorphisms | 3696 | - | - |
| **Mean nucleotide diversity estimates\*** |  |  |  |
| SNPs only | 3.849E-03 | **3.957E-03\*\*** | **3.740E-03** |
| " " diversity range |  | 0-0.03823 | 0-0.02525 |
| Tajima's evolutionary standard deviation | 2.549E-03 | 2.632E-03 | 2.465E-03 |
| SNPs only (509 chosen genes) | 3.752E-03 | 3.821E-03 | 3.682E-03 |
| SNPs only (202 random genes) | 4.103E-03 | **4.306E-03** | **3.900E-03** |
| All polymorphisms | 4.359E-03 | **4.471E-03** | **4.247E-03** |
| " " diversity range |  | 0-0.03893 | 0-0.02525 |
| Tajima's evolutionary standard deviation | 2.816E-03 | 2.903E-03 | 2.729E-03 |
| All polymorphisms (509 chosen genes) | 4.214E-03 | 4.278E-03 | 4.150E-03 |
| All polymorphisms (202 random genes) | 4.716E-03 | **4.944E-03** | **4.488E-03** |

Also, 48% and 52% of the polymorphisms observed were exclusive to *Q. petraea* and *Q. robur* respectively in our panel, the rest being shared among species (Table 3). Among exclusive polymorphisms, 46% and 44% were singletons in *Q. petraea* and *Q. robur* respectively, suggesting that they might be either rare in both species, or more polymorphic in local populations from which few individuals were sampled across the species wider ranges. Overall and within both species, we observed a large variation in the numbers of segregating sites per gene size (Fig. S7-B, Supporting information).

The mean nucleotide diversity estimates (*θπ*) across genic regions when considering all polymorphisms were 0.00447 and 0.00425 in *Q. petraea* and *Q. robur* respectively, with up to a 10-fold variation among polymorphic genes overall and in different functional categories (Fig. 4-B and Table 3).

When including SNPs only, mean *θπ* decreased overall by more than 10% (Table 3, and see column D in Table S4, Supporting information). The large variation among genes is also illustrated by the absence of significant differences between mean diversity among functional categories *within species*, in most comparisons using non-parametric Wilcoxon rank sum tests (*Wrs*) with similar number of genes. Two notable exceptions were observed when considering all polymorphisms: the *biotic stress* category (358 genes) had on average a lower *θπ* in *Q. petraea* than in the random gene list (211 genes, *Wrs* Pr<0.042), and the mean *θπ* of the *reproductive phenology* category was significantly lower in both species than that of the *Bud phenology* category (*Wrs* Pr<0.040 and Pr<0.013 in *Q. petraea* and *Q. robur* respectively, considering exclusive categories from Table S2, Supporting information). Genes with *θπ* estimates above 0.02 were found across most categories, whether considering all polymorphisms (Fig. 4-B) or SNPs only. The 8 genic regions showing the highest *θπ* values in both species were annotated for example as disease resistance, transcription factor or membrane transport proteins, half of them being from the original random list.

Comparing nucleotide diversity between individuals according to their main cpDNA lineages B *versus* A or C (Table 1), no significant differences were found between lineages within both species, using *Wpr* tests across all genes (see also the lineage-associated distributions of genes’ diversity in Fig. S8, Supporting information). This was also true for all functional categories. In both species, the mean differentiation across genes among lineages was very low (<0.015, each gene estimate being the mean *F*_*ST*_ across all polymorphisms at this gene), with very few genes (~1%) having much higher mean *F*_*ST*_ (ranging from 0.21 to 0.41 or 0.56 within *Q. petraea* and *Q. robur* respectively).

Mean *θπ* comparison tests *between species* across all gene regions were not significant (Table 3, *Wrs* Pr>0.15 for all polymorphisms or SNPs only), nor were they across different categories and between gene pairs, using a 95% confidence interval based on Tajima’s evolutionary variance for *θπ* (Tajima 1983) while assuming underlying Gaussian distributions. Indeed for the same genic regions, many examples can be found of higher *θπ* estimates in one species or the other. However, comparing diversity estimates across the exact same positions and performing Wilcoxon paired ranked tests (*Wpr*) across all genes, there was a significant pattern of a slightly higher diversity in *Q. petraea* (see Table 3 and Fig. 4-B), whether considering all polymorphisms (*Wpr* Pr<0.028) or SNPs only (*Wpr* Pr<0.036). This pattern remained significant across the 202 polymorphic genes chosen randomly (*Wpr* Pr<0.037, all polymorphisms, Table 3), even when excluding the 5% or 10% of genes having the highest *θπ* values. This pattern of a significantly higher *θπ* in *Q. petraea* was not observed when considering the 509 polymorphic gene regions chosen in functional categories, either together or separately in the different categories (Fig. 4-B), except for the *Abiotic stress* category.

We also observed a very large variation for *F*_*ST*_ estimates across gene regions and functional categories, which covered the full range of possible values [0,1], with mean values of ~0.13 whether considering all polymorphisms or SNPs only (Fig. 4-C, and Fig. 4-D for the random genic regions and a representative example in one category). The very few segregating sites with *F*_*ST*_ values equal to one had either missing individuals’ or strands, possibly caused by polymorphisms within primer regions. Among the sites sequenced for the full sample of gametes, the 20 highest *F*_*ST*_ values ranged from 0.6 to 0.9 and belonged to 10 genic regions, many of which also showed null or very low *F*_*ST*_ values within 100 bp. This large variation in differentiation was observed between very close variant sites in many genes, suggesting very high recombination rates at genome-wide and range-wide scales, and consistently with the very low expected background LD (see above). Additionally, a large variance is expected around *F*_*ST*_ estimates due to the relatively low sample size in both species, in particular for bi-allelic loci (Weir and Hill 2002; Buerkle *et al*. 2011; e.g. Eveno *et al.* 2008).

## Discussion

In the NGS era, non-model tree species such as many *Fagaceae* still lag behind model species for easy access to sequence polymorphism and SNP data (but see Gugger *et al*. 2016 for *Quercus lobata*). These data are needed for larger scale studies addressing the many diversity issues raised by their combined economic, ecological and conservation interests (Cavender-Bares 2016; Fetter *et al*. 2017; Holliday *et al*. 2017). Recent achievements and data availability from the *Q. robur* genome sequence project (Plomion *et al*. 2018) opens a large range of applications in many related temperate and tropical *Fagaceae* species due to their conserved synteny (Cannon *et al.* 2018). In this context, we discuss below the representativity of our data in terms of species genomic diversity as well as the robust patterns observed across genes, and further illustrate their past and future usefulness for *Quercus* species.

### Genic resources content, quality, and representativity

We provide a high-quality polymorphism catalog based on Sanger resequencing data for more than 850 gene regions covering ~530 kb, using a discovery panel (*DiP*) from mixed *Q. robur* and *Q. petraea* populations located in the western and central European part of their geographic range. This catalog details functional annotations, previous published information, allele types, frequencies and various summary statistics within and across species, which can assist in choosing novel polymorphic sites (SNPs, SSRs, indels…) for genotyping studies. Among genomic SSRs, more than 90% (~200) are new (17 already detected in Durand *et al*. 2010; 3 in Guichoux *et al.* 2011), so they constitute an easy source of potentially polymorphic markers in these oak species. Standard formats for high-density genotyping arrays and primer information are also provided, making these resources readily operational for medium scale molecular ecology studies while avoiding the burden of bioinformatics work needed for SNP development (Tables S1 to S5, Supporting information, and see also https://github.com/garniergere/Reference.Db.SNPs.Quercus for additional information). This catalog corrects and largely extends the SNP database for Q. petraea/robur at https://arachne.pierroton.inra.fr/QuercusPortal/ which was previously used to document a SNP diversity surrogate for both *Quercus* species in the oak genome first public release (Plomion *et al*. 2016).

Thanks to a high quality dedicated pipeline, we could perform a quasi-exhaustive characterization of polymorphism types in our *DiP* and across part of the genic partition of these *Quercus* species (see Fig. 1). Although base call error rates below 1/1000 were used (as originally developed for Sanger sequencing), most variant sites were located in regions with lower error rates (below 1/10000) so that true singletons could be identified. At the genotypic level, a Sanger genotyping error rate below 1% was previously estimated using a preliminary subset of around 1200 SNPs from this catalog (corresponding to around 5800 data points in Lepoittevin *et al.* 2015). This rate can be considered as an upper bound for the present study, given all additional validation and error correction steps performed. Although little produced now with the advent of NGS methods, Sanger data have served for genome sequencing projects in tree species before 2010 (Neale *et al*. 2017), and have been instrumental, in combination to NGS for BAC clones sequencing, in ensuring assembly long-distance contiguity in large genomes such as oaks (Faivre-Rampant *et al*. 2011, Plomion *et al*. 2016). Sanger sequencing has also provided reference high-quality data to estimate false discovery or error rates, and validate putative SNPs in larger scale projects (e.g. Geraldes *et al*. 2011 in *Populus trichocarpa*; Sonah *et al*. 2013 in Soybean; Cao *et al*. 2014 in *Prunus persica*).

Finding an optimal balance between the number of samples and that of loci is critical when aiming to provide accurate estimates of diversity or differentiation in population genetics studies. Given the increasing availability of markers in non-model species (usually SNPs), it has been shown by simulation (Willing *et al*. 2012, Hivert *et al*. 2018) and empirical data (Nazareno *et al*. 2017) that sample sizes as small as 4 to 6 individuals can be sufficient to infer differentiation when a large number of bi-allelic loci (> 1000) are being used. A broad-scale geographic sampling is however required if the aim is to better infer genetic structure and complex demographic scenarios involving recolonization and range shifts due to past glacial cycles, such as those assumed for many European species (Lascoux and Petit 2010, Keller *et al*. 2010, Jeffries *et al*. 2016, Sousa *et al*. 2014). Our sampling design is likely to have targeted a large part of both species overall diversity and differentiation across the resequenced genic regions. This is first suggested by the small proportion of additional polymorphisms once an initial sample of 8 gametes was included for each species (i.e. ~10% and decreasing as sample size increases, Fig. 4-A and Fig. S7-A, Supporting information). Considering the *DiP* within each species, each individual brings on average ~166 new variants (~1% of the total). Second, the large variance observed across gene nucleotide diversity estimates (see Table 3) is mostly due to stochastic evolutionary factors rather than to sampling effects so unlikely to be impacted by sample sizes over 10 gametes (Tajima 1983). Third, sampling sites are located in regions which include 4 out of the 5 main cpDNA lineages reflecting white oaks recolonization routes (lineages A to C and E in Petit *et al.* 2002a), the likely haplotypes carried by the *DiP* individuals being A to C (Table 1).

Therefore, if new populations were being sampled within the geographical range considered, they would likely include many of the alleles observed here within species and at other genes across their genomes. For differentiation patterns, older and more recent reports showed a low genetic structure among distant populations within each species, and a relatively stable overall differentiation among species compared to possible variation across geographical regions (Bodénès *et al.* 1997; Mariette *et al*. 2002; Petit *et al*. 2003; Muir and Schlötterer 2005; Derory *et al.* 2010; Guichoux *et al*. 2013; Gerber *et al*. 2014). For new populations sampled outside the *DiP* geographic range, a recent application to *Q. robur* provenances located in the low-latitude range margins of the distribution (where 3 main cpDNA lineages occur) showed a high rate of genotyping success, a high SNP diversity, and outliers potentially involved in abiotic stress response (Temunovic *et al*. 2020).

We further tested the frequency spectrum representativity of our range-wide *DiP* by comparing genotypic data for a set of 530 independent SNPs (called *sanSNP* for Sanger data) with data for the same set of SNPs obtained in Lepoittevin *et al*. (2015, called the *illuSNP* set since it used the Illumina infinium array technology) for larger numbers of ~70 individuals per species from Southern France natural stands. The SNPs were chosen so that the *illuSNP* set excluded SNPs showing compressed clusters (*i.e*. potential paralogs) and those showing a high number of inconsistencies with control genotypes, as recommended by the authors. Comparing between datasets, for SNPs exclusive to one species in the *sanSNP* set, more than 68% either show the same pattern in the *illuSNP* set, or one where the alternative allele was at a frequency below 5% in the other species. Less than 8% of those SNPs are common in both species in the *illuSNP* set. Similarly, for singletons in the *sanSNP* set, more than two-third of the corresponding SNPs in the *illuSNP* set showed very low to low frequency (<10%), while only 11% in *Q. petraea* and 9% in *Q. robur* showed a *maf* above 0.25. This further confirms the reality of singletons in our *DiP*, and also that some may represent more frequent polymorphisms in larger samples of local populations. The correlations among *maf* in both datasets were high and significant (0.66 and 0.68 respectively for *Q. petraea* and *Q. robur*, both Pr< 0.0001).

Finally, various methodological steps and obtained results tend to demonstrate that we avoided a bias towards low-diversity genic regions: (i) an initial verification that very low BlastX *E*-values (< 10^−80^) did not target more conserved regions, (ii) a primer design optimizing the amplification of polymorphic fragments, both (i) and (ii) using potential variants in ESTs data assembled across both species (Fig. S2-B steps 1 and 3; Appendix S1, Supporting information), (iii) a high nucleotide diversity across genes and ~50% of shared variants (Table 3 and Fig. 4), (iv) a very low proportion of fragments with no detected variants, and a substantial part (~30%) of variant positions due to Indels and SSRs (Table 2), (v) additional results showing that, across ~100 kb of more than 150 independent fragments amplifying in one species only and thus with possible more divergent primer pairs, the number of detected heterozygotes was twice smaller compared to fragments amplifying in both species (more details in Appendix S1, Supporting information).

These results altogether suggest a small risk of SNP ascertainment bias if these new resources were to be used in populations both within and/or outside the geographic distribution surveyed, in contrast to panels with much less individuals than here (see respectively Lepoittevin *et al*. 2015 for a discussion on the consequences of such bias in *Quercus* species, and Temunovic *et al*. 2020 cited above).

Overall, we obtained sequence data for 0.072% (~530 kb) of the haploid genome of *Q. robur* (size of ~740 Mb in Kremer *et al*. 2007). We also targeted ~3% of the 25808 gene models described in the oak genome sequencing project (www.oakgenome.fr), and around 1% of the gene space in length. Interestingly, both randomly chosen genic regions and those covering different functional categories have been mapped across all linkage groups (columns F and X in Table S1, Supporting information). Due to the absence of observed background LD, their diversity patterns can be considered independent. The genes studied represent a large number of categories, as illustrated by very similar distributions for level 2 GO terms to those obtained with the larger *ocv4* assembly (Lesur *et al*. 2015, comparing their Figure 2 to Fig. S4-A to S4-C, Supporting information).

### Diversity magnitude and heterogeneity highlight species integrity and introgression patterns

Using a detailed polymorphism typology, we characterized for the first time in two oak species a high proportion of variant positions (30%) that included one bp to medium-sized indels and sequence repeats, compared to the more common and commonly reported SNP loci (Table 2). The proportions of indels observed (11.5% of all polymorphisms) is in the range of results available in model tree species (e.g. 13.8% across the genome in *Prunus avium*, Shirasawa *et al*. 2017; 19% in *Prunus persica,* Cao *et al*. 2014; a lower estimate of 1.4% in *Populus trichocarpa*, Evans *et al*. 2014). Although less abundant than SNPs, they represent an important component of nucleotide variation, often having high functional impacts when located within coding sequences, and they have been proposed as an easy source of markers for natural populations studies (Väli *et al*. 2008). Larger-sized indels are also likely to be relatively frequent in intergenic regions of the *Quercus* genome and have been linked to transposable elements (TE, see the BAC clones overlapping regions analyses in Plomion *et al*. 2016). Similarly, large indels and copy number variation linked to TE activity were identified as an important component of variation among hybridizing *Populus* species (Pinosio *et al*. 2017). Here when considering variant positions involved in complex polymorphisms, we observed one variant position per 48 bp on average within species (resp. one per 30 bp in both), compared to the one SNP per 68 bp statistic (resp. one SNP per 42 bp across both species). Also, some of the SNPs observed were located within complex polymorphic regions that would have been classically filtered out, and nucleotide diversity (*θπ*) estimates were higher by 12% when including all polymorphisms (from 0.0038 to 0.0044 if averaging across both species and all genes, Table 3). These *θπ* estimates are provided for the first time in *Q. petraea* and *Q. robur* across a large number genic regions (> 850), compared to previous candidate genes studies across much smaller numbers (< 10) of gene fragments (Kremer *et al.* 2012 in *Q. petraea*; e.g. Homolka *et al*. 2013).

Based on these data, there is an interest in attempting to estimate SNP numbers across the full genome of the studied species for range-wide samples, as it may impact filtering strategies in pipelines for future NGS haplotype-based data production, or decisions to develop or not SNP arrays in these species. In order to do that, a few realistic assumptions can be made from both the exhaustive description of variants provided, and the mean proportions of SNP numbers in new individuals that we computed for increasing across sample sizes. First ~10% additional rare SNPs per sample could be observed for a *DiP* twice as large as ours (based on Fig. S7-A data, Supporting information). Thus given the representativity of our data compared to the *ocv4* unigene (Lesur *et al.* 2015), we would expect around 1.36 million SNPs on average within species by applying our statistics to the full genic partition of *Q. robur* or *Q. petraea* (~80 Mb, www.oakgenome.fr, Plomion *et al.* 2018). Another reasonable assumption is that shared and exclusive polymorphisms proportions across genic regions would be around 30% and 70% respectively, for these closely related oak species (based on both our *DiP* and Lepoittevin *et al*. 2015 results), which translates into the presence of ~2.32 million SNPs for the genic partition in a sample including both *Q. petraea* and *Q. robur* (resp. ~4.22 if including also *Q. pubescens* and *Q. pyrenaica*). Finally, if we apply to the *Quercus* genome a range of ratios for SNPs counts in intergenic over genic regions estimated from several tree species natural population samples (2.03 in *Populus trichocarpa*, Zhou and Holliday 2012; 2.25 in the “3P” *Q. robur* reference genotype, Plomion *et al.* 2016; 2.57 in *Prunus persica* wild accessions, Cao *et al.* 2014), we obtain an estimate of between 34 to 42 million SNPs within species across a large spatial range (resp. 41 to 51 million SNPs in both Q. petraea and robur species, and 75 to 94 million SNPs considering the 4 species previously cited). All these figures could be at least 30% higher if one considers all possible variants involved in indels, SSRs and complex polymorphisms, as shown in our results. Although of the same order of magnitude, the contrast with the twice smaller number of SNPs identified in Leroy *et al.* 2019 (~32 millions) across the same four species with similar sample sizes than ours, could be explained by different factors. First their filtering strategy applied on Pool-seq data in order to minimize errors basically excludes all singletons. However, we have seen that verified singletons which could represent rare or local variants amounted to more than 20% of all polymorphisms (see Results). Indeed, very stringent filters are often applied in practice to limit error rates and avoid false-positives, hence limiting the impact of variable read depth and possible ascertainment bias risks, which altogether significantly decrease the number of informative loci compared to either initial fixed amounts (in genotyping arrays, e.g. Lepoittevin *et al.* 2015) or potential amounts (in reference genomes, e.g. Pina-Martins *et al*. 2019 in *Quercus* species; see also Van Dijk *et al*. 2014). Second, no cross-validation step is available in Leroy *et al.* (2019) for data quality, that would have permitted to have a better grasp of possible bias and error rate expected in such a dataset, and its consequences on allele frequency estimates and inference methods (see Hivert *et al*. 2018 and discussion below). Also, we can’t exclude that a regional sampling strategy such as the one used in Leroy *et al*. (2019) might miss allelic variants with a higher *maf* in other regions for the two species having the wider geographical range.

Our *θπ* estimates are consistent with those obtained from genome-wide data and range-wide panels in angiosperm tree species, available mostly from the model genus *Populus* (e.g. *P. trichocarpa*: one SNP per 52 bp and *θπ*~0.003 across genic regions, Zhou and Holliday 2012, Zhou *et al.* 2014, Evans *et al*. 2014, Wang *et al*. 2016; *P. tremula*: *θπ*~0.008, P. tremuloides: *θπ*~0.009 across genic regions, Wang *et al*. 2016; *θπ*~0.0026 to 0.0045 in a panel including wild *Prunus persica* accessions, Cao *et al.* 2014). These diversity levels are also within the range estimated for the long-term perennial outcrosser category in Chen *et al*. (2017, see Fig. 1-D with a mean value of silent *θπ*close to ~0.005) and can be considered relatively high in the plant kingdom if excluding annual outcrosser estimates or intermediate otherwise. In oaks as in many other tree species with similar life history traits, these high levels would be consistent with their longevity, large variance in reproductive success and recolonization or introgression histories, which could have maintained deleterious loads of various origins (Zhang *et al*. 2016, Chen *et al*. 2017, Christe *et al.* 2016b).

Comparing the nucleotide diversity distributions and examining the range of differentiation across genic regions in our *Dip* reveal several robust patterns that altogether illustrate historical introgression among both *Quercus* species. These two species have long been considered as iconic examples of species exhibiting high levels of gene flow (e.g: Petit *et al*. 2003; Arnold 2006), despite more recent evidence of strong reproductive barriers (Abadie *et al.* 2012). What has been referred to as “strong species integration” seems nevertheless clearer in our *Dip* for *Q. robur* than for *Q. petraea*, according to genetic clustering inference without any *a priori*. Three individuals (27%) considered as typical morphological *Q. petraea* adults (Kremer *et al* 2002a) showed significant levels of introgression (Fig. 3). In contrast, only one *Q. robur* based on morphology was introgressed to a level matching the least introgressed *Q. petraea* individual. Discussing species delimitation, Guichoux *et al*. (2013) also showed more robustness in assigning morphological *Q. robur* individuals to their genetic cluster, illustrating an asymmetry in their introgression levels. We note that among our *Dip* individuals, Qs28, one parent from two mapping pedigrees (Bodénès *et al*. 2016) is a clear F1 hybrid among both species (Fig. 3), making those pedigrees two back-crosses instead of one cross within species and one between species.

Moreover, after excluding the four most introgressed individuals, nucleotide diversity in *Q. petraea* was significantly higher (by ~5% on average) than in *Q. robur*. This effect is small, detectable only with Wilcoxon paired ranked tests, mostly across the same ~200 regions sampled randomly and in the *Abiotic stress* category, despite the very large diversity variance across regions, and robust to excluding the highest diversity values. We also sequentially removed the three individuals with the highest *Q*-values from the *Q. petraea* cluster (Fig. 3), since they could still harbor residual heterozygosity due to recent back-crossing events and generate the pattern observed. Remarkably, the same significant patterns of higher diversity in *Q. petraea* were observed. Therefore, with 8 to 10 gametes in *Q. petraea* instead of 8 to 24 gametes in *Q. robur*, and with twice less natural stands sampled, the nucleotide diversity in *Q. petraea* was still slightly and significantly higher than in *Q. robur* (Pr<0.011 and Pr<0.026, using all polymorphisms or SNPs only respectively). Although the magnitudes of range-wide population structure within both species could differentially affect both species global diversity across our *Dip*, published results show that these are very small with similar values (~1% across SNPs, Guichoux *et al.* 2013).

The main hypotheses proposed so far to explain this difference in extent of diversity between species relate to their disparities in life-history strategies for colonizing new stands and associated predictions (Petit *et al*. 2003, Guichoux *et al*. 2013). The colonization dynamics model and patterns observed also assumes very similar effective population sizes in both species, which is a reasonable assumption due to their shared past history and the strong introgression impact at the genomic level. However, given increasing and recent evidence of pervasive effects of different types of selection across genic regions with high-throughput data (e.g. Zhang *et al*. 2016; Christe *et al.* 2016b in *Populus*; Chen *et al* 2017 for long-term perennials), alternative (and non-exclusive) hypotheses worth considering are ones of a higher genome-wide impact of selective constraints in *Q. robur (*Gillespie 2000; Hahn 2008; Cutter and Payseur 2013; Kern and Hahn 2018; e.g. Grivet *et al.* 2017). Since *Q. robur* is the most pioneering species, it has likely been submitted to very strong environmental pressures at the time of stand establishment. Selection might be efficient, given oak tree reproductive capacities, and affect variation across a large number of genes involved in abiotic and biotic responses. This would be consistent with significantly lower levels of diversity (*He*) in *Q. robur* at SNPs located in genes that were specifically enriched for abiotic stress GO terms (Guichoux *et al*. 2013, see their Table S5). Redoing here the same tests across a larger number of independent SNPs (> 1000), *Q. petraea* systematically showed the same trend of a slightly higher diversity overall, and significantly so only for the *Abiotic stress* category (*Pr*<0.01) and for a similar outlier SNP category (*F*_*ST*_ >0.4, mean He>0.15, *Pr*<0.001) than in Guichoux *et al*. (2013). In summary, the absence of the same pattern in any other functional categories might suggest that these are too broad in terms of corresponding biological pathways, hence mixing possible selection signals of opposite effects among species, while we still detect an overall effect due to linked selection on a random set of genes, and on genes involved in abiotic stress.

Within both species, no differences in nucleotide diversity, and a very small differentiation (below 1.5%) were found on average across genes among the main cpDNA lineages (B *versus* A or C) that indicate past refugial areas and migration routes. These patterns were expected, given oaks’ life history traits (e.g. high fecundity and dispersal rates), large population sizes, and plausible recolonization scenarios throughout Europe leading to current levels of adaptive differentiation among populations at both nuclear genes and traits (Kremer *et al*. 2010). Only cpDNA ancient differentiation signals among isolated historical refugia were retained, while other putative adaptive divergence effects due to different environments were erased, as illustrated by an absence of correlations between cpDNA and nuclear or phenotypic traits divergence across populations (Kremer *et al*. 2002b). This is consistent with many events of population admixture during the last ~6000 thousands years after European regions were recolonized, as well as a very low genetic differentiation among distant populations (e.g. Guichoux *et al*. 2013), which contrasts with a much higher differentiation often observed for adaptive traits (e.g. Kremer *et al*. 2014; Sáenz-Romero *et al*. 2017). Interestingly, the very few genes with mean *F*_*ST*_ between 0.21 and 0.56 among lineages are not the same in *Q. petraea* and *Q. robur* (five and seven genes respectively). Seven of them have GO terms indicating their likely expression in chloroplasts, or their interaction with chloroplastic functions. They are either housekeeping genes for basic cellular functions, or belong to biotic or abiotic stress functions (seven of them), and could be involved in local adaptation between ecologically distant populations, calling for further research in larger samples.

More generally, analyses comparing the nucleotide diversity patterns at genes involved in both species relevant biosynthesis pathways for ecological preferences (e.g. Porth *et al*. 2005; Le Provost *et al*. 2012, 2016) are clearly needed in replicated populations, for example to estimate the distribution and direction of selection effects and putative fitness impact across polymorphic sites (Stoletzki and EyreWalker 2011), or to study the interplay between different types of selection and variation in local recombination rates on both diversity and differentiation patterns (Payseur and Rieseberg 2016).

A large proportion of shared polymorphic sites (~50% in any species) highlights the close proximity of species at the genomic level, consistently with a low mean differentiation across polymorphic sites (*F*_*ST*_~0.13, Fig. 4-C), and despite the very large heterogeneity observed across differentiation estimates. This has now been classically interpreted (and modeled) as reflecting a strong variance in migration and introgression rates, in oaks in particular (Leroy *et al*. 2017), with islands of differentiation assumed to represent regions resistant to introgression. However, interpretations of such patterns remain controversial and multiple processes might be involved and worth exploring further in oaks, such as the effects of heterogeneous selection (both positive and background) at linked loci (Cruickshank and Hahn 2014; Wolf and Ellegren 2017). These effects could be particularly visible in low-recombination regions (Ortiz-Barrientos *et al*. 2016), and would further interact with the mutational and recombination landscapes during the course of speciation (Ortiz-Barrientos and James 2017) and during their complex demographic history.

### Applications and usefulness as reference data

During this project, several studies valued part of these resources, hence illustrating their usefulness. For example, good quality homologous sequences were also obtained for ~50 % of the gene fragments in one individual of *Quercus ilex*. This species is relatively distant genetically to both *Q. petraea* and *Q. robur*, belonging to a different section, so these data guided the choice of nuclear genes for better inferring phylogenetic relationships across 108 oak species (Hubert *et al.* 2014). Bioinformatics tools and candidate genes annotated during the project were also useful to similar genes and SNP discovery approach in *Quercus* or more distant *Fagaceae* species (Rellstab *et al*. 2016, Lalagüe *et al.* 2014 in *Fagus sylvatica,* El Mujtar *et al*. 2014 in *Nothofagus* species). Given the low ascertainment bias and good conversion rate expected within the range surveyed, those genomic resources would be directly applicable to landscape genomics studies at various spatial scales (reviewed in Fetter *et al.* 2017) in both *Quercus* species. Indeed, easy filtering on provided SNP statistics in the catalog would allow distinguishing among different classes of SNPs (e.g. exclusive to each species, common and shared by both, linked to particular GO functional categories), delimiting and tracing species in parentage analyses and conservation studies (e.g. Guichoux *et al*. 2013; Blanc-Jolivet *et al.* 2015), or improving estimates of lifetime reproductive success and aiming to understand how demographic history and ecological drivers of selection affect spatial patterns of diversity or isolating barriers (Andrew *et al*. 2013; e.g. Geraldes *et al*. 2014). This type of spatial studies are surprisingly rare in these oak species, they usually include a small number of SSR markers, and all suggest complexity in geographical patterns of genetic variation and importance of the ecological context (e.g Neophytou *et al*. 2010; Lagache *et al*. 2014; Klein *et al*. 2017, Beatty *et al.* 2016 for local or regional studies; Muir and Schlötterer 2005; Gerber *et al*. 2014, Porth *et al.* 2016 for range-wide studies). Their power and scope would likely be greatly improved by using medium-scale genotyping dataset including a few thousands SNPs such as those described in our study.

The robust patterns described above of differentiation heterogeneity and consistent differences in diversity magnitude among species call for more studies at both spatial and genomic scales for unraveling these species evolutionary history, in particular regarding the timing, tempo, dynamics and genetic basis of divergence and introgression. Practically, in order to address those questions in oaks, genomic data on larger samples of individuals could be obtained from either genome complexity reduction methods such as RAD-seq and similar approaches (e.g. Andrews *et al.* 2016) or previously developed SNP arrays (e.g. Silva-Junior-*et al*. 2015). We do not recommend the development of a very large SNP array in oaks since it is likely to be very costly for the actual return, especially given the very large and range-wide panel that would be needed to significantly limit ascertainment bias (see Lepoittevin *et al*. 2015). The very low overall levels of LD observed here indicate also potentially high recombination rates, and thus that a very high SNP density would be required for targeting functional variants, which would not be compatible with technical constraints for controlling for genotyping error rates (previously shown to be high in SNP array). Indeed, these rates would probably be stronger for high diversity, complex, duplicate or multiple copy genic regions (as those observed in this study in Tables S1 and S4, Supporting information, and shown recently to have an evolutionary impact on the *Q. robur* genome structure, Plomion *et al*. 2018), preventing these regions to be included in SNP arrays. The very short LD blocks observed in this study might also limit the utility of RADseq data alone to uncover many loci potentially under selection in genome scans for local adaptation studies (Lowry *et al.* 2016; McKinney *et al*. 2017). In contrast, targeted sequence capture (TSC) strategies for resequencing (Jones and Good 2016), and the more recent advances in RADseq approaches that deal with previous limitations (Arnold *et al*. 2013; Henning *et al*. 2014; and see Rochette *et al*. 2019), although still uncommon in forest tree species evolutionary studies, might be more useful and efficient since they can be oriented towards recovering long genomic fragments. They would thus allow more powerful site frequency spectrum and haplotype-based inferences to be pursued, therefore avoiding most of the SNP arrays technical issues (e.g. Zhou *et al.* 2014; Wang *et al*. 2016), especially given the large variance in nucleotide diversity and low overall differentiation characterized here. TSC approaches will surely be encouraged and tailored to specific evolutionary research questions in oaks in the next decade, given the new *Q. robur* genome sequence availability (Plomion *et al*. 2018; Lesur *et al*. 2018 for the first TSC in oaks). However, the bioinformatics pipelines needed for validating haplotype-based or quality data for population genetics inferences also need constant reassessment according to research questions and chosen technology.

We thus propose, in addition to direct applications to landscape genetics (detailed above) and transferability to other *Quercus* species (for example using primer information in Table S1, Supporting information, and see Chen *et al*. 2016), that the high-quality data characterized in this study serve as a reference for such validation purposes. They could not only help for adjusting parameters in pipelines for data outputs, but also allow estimating genotyping error rates for SNP and more complex classes of variants, either by comparing general patterns (e.g *maf* distribution from Tables S3, S4 Supporting information) or using the same control individuals maintained in common garden that could be included in larger-scale studies. Such a reference catalog of SNPs and other types of polymorphisms within gene fragments could also be very useful for solid cross-validation of variants identification, allele frequency and other derived summary statistics in alternative strategies such as *Pool-Seq*, which allow increasing genomic coverage while sampling cost-effectively by pooling individuals (Schlötterer *et al.* 2014). Indeed, the drawback of *Pool-Seq* approaches, despite dedicated software (PoPoolation2, Kofler *et al*. 2011) is that they can give strongly biaised estimates, or ones that do not consider evolutionary sampling (Hivert *et al*. 2018). Therefore, they require further validation methods which usually value previously developed high-quality and lower-scale data (e.g. Pool-Seq *versus* Sanger and Rad-Seq in Christe *et al*. 2016b; Illumina GA2 *versus* Sanger in Cao *et al*. 2014; EUChip60K *versus* deep-whole genome resequencing in Silva-Junior *et al.* 2015). Finally such a reference dataset would help optimizing the amount of data recovery from either TSC or whole-genome resequencing experiments in future research challenges by fine-tuning dedicated data processing bioinformatics pipelines.

## Author contributions

Funding acquisition: AK, PGG, CP, and MLDL; Initial conception and individuals sampling: PGG, AK, CP, MPR, VL; Bioinformatics strategy and experimental design: PGG, TL; DNA extraction and quality check: VL; Sequence Data acquisition: PGG, CP, TL, VL; Individuals’ identification checks for quality control VL, CL, PL; Pilot study: VL, PGG; Working assembly: JMF, PGG: Primer design and amplicon choice: PGG, VL, TD; Original candidate gene lists choice: PGG, TL, JMF, CP, AK, TD, CR, MLDL, GLP, ChB, EG, CaB, NT, PA; Bioinformatics tools: TL and PGG (SeqQual pipeline and R scripts), JMF and AF (Bioperl and R scripts), PA, CL, VelM, JT, FH, TD (SeqQual tests), FR (website); Visual Chromatogram checks, SNP/assembly validations: PGG, VL, TD, PA, TL, MLDL, CaB, ChB, CL, CR and EG; Bioinformatic and population genetic analyses: PGG, TL, SM, ChB: Functional annotation: TL, PGG, VelM, PA; Manuscript draft: PGG; Manuscript review and edition: PGG, SM, CL, ChB, TL; all authors agreed on the manuscript.

## Data and code Accessibility

The original assembly used for selecting contigs is in Appendix S2 (Supporting information). For Sanger trace files (with data on at least 2 individuals), see the Dryad repository (at the https://doi.org/10.5061/dryad.4mw6m906j link). Consensus sequences are respectively in appendices S3 (used to design primers and for functional annotation, see also Table S2), S4 (genomic sequences obtained), and S5 (genomic sequences obtained for *Q. ilex*). Tables S1 and S2 correct and extend the current *Q. robur and Q. petraea* Candidate Genes Database of the Quercus Portal (www.evoltree.eu/index.php/e-recources/databases/candidate-genes, accessible by selecting “Fagaceae” then “All” in the “Refine Data” Menu). SNP, indel and SSR catalogs and positions within genomic consensus sequences, and ready-to-use format for genotyping essays are provided in Tables S3 to S5 (Supporting information), and at https://github.com/garniergere/Reference.Db.SNPs.Quercus with additional information.

Bioperl scripts from the SeqQual pipeline are given at https://github.com/garniergere/SeqQual, example of parameter files and scripts for STRUCTURE analyses and parsing MREPS software are given at https://github.com/garniergere/Reference.Db.SNPs.Quercus

## Acknowledgments

The authors thank Alexis Ducousso, Jean-Marc Louvet, Guy Roussel, Pablo Goicoecha, Hervé le Bouler, Félix Gugerli, Csaba Matyas, Sandor Bordacs, Hans P. Koelewijn, Joukje Buiteveld, Stephen Cavers, Bernd Degen and Jutta Buschbom for choosing trees and providing dried leaves of individuals from various Intensive Study Populations of previous European projects populations. We are grateful to H. Lalagüe, G. Vendramin, I. Scotti, and L. Brousseau for testing earlier scripts of SeqQual and to I. Lesur for help in using the *ocv4* oak resources. The sequencing work was funded by the EVOLTREE network of Excellence (EU contract n°016322). TL post-doc fellowship was funded by the ANR TRANSBIODIV (06-BDIV-003-04) and LINKTREE (contract n°2008-966). TD salary was funded by the ANR REALTIME (N°59000256). Computing facilities of the Mésocentre de calcul Intensif Aquitain des Universités de Bordeaux, de Pau et des Pays de l’Adour are thanked for providing computer time for this study. We also thank Rémy Petit for funding part of TL fellowship and support in developing SeqQual tools. PA received a Ph.D. grant (2009-2011) from the « Ministère de l’Education Nationale, de l’Enseignement Supérieur et de la Recherche » of France, and additional funding from EVOLTREE. We also gratefully thank Oliver Brendel, Ricardo Alia, Komlan Avia and Hilke Schröder for their time and constructive comments. Version 4 of this preprint has been peer-reviewed and recommended by Peer Community In Forest and Wood Sciences (https://doi.org/10.24072/pci.forestwoodsci.100003).

## Conflict of interest disclosure

The authors of this article declare that they have no financial conflict of interest with the content of this article.

## Supporting information

All supporting information is available in the BioRxiv server: https://doi.org/10.1101/388447

**Fig. S1** Sampling site locations within the natural geographic distribution of *Q. petraea* and *Q. robur*. Vector map is from *http://www.naturalearthdata.com* and distribution areas from Euforgen (*http://www.euforgen.org/distribution-maps/)*

**Fig. S2** Working assembly steps and softwares (A), and bioinformatic strategy for search of candidate genes and amplicon choice (B).

**Fig. S3** Plots of the ΔK values from the Evanno *et al*. (2005) method (S3-A, -B, -C, -D, -E), and of the mean values of the estimated probability ln (of the data given K) with standard deviations for K ranging from 1 to 5 (S3-F to S3-J), which show support for K=2. Plots are from the STRUCTURE HARVESTER program.

**Fig. S4** Distributions of Gene Ontology (GO) terms for the consensus sequences in Appendix S3, at level 2 (-A,-B, -C) and level 3 (-D, -E, -F): A- and D- for Biological Process, B- and E- for Molecular Function, C- and F- for Cellular Component. Annotation rules: E-value<10^−30^, annotation cut-off 70, GO weight 5, HSP coverage cutoff 33%. Filtering applies for at least 5 sequences and a node score of 5 per GO term (but see rare exceptions in Table S2).

**Fig. S5** Distributions of GO terms across different gene lists (*bud*, *abiotic* and *biotic*) at Biological Level 2, and Fisher exact tests across pairs of sequence clusters with the same GO terms between the random list and other lists. Significance levels *: P<0.05.

**Fig. S6-A to S6-J** Posterior assignment probabilities (*Q-*values) of 24 individuals attributed to 2 clusters (STRUCTURE analysis) for different numbers of polymorphisms, different sampling of SNP data, and different plots of credible intervals.

**Fig. S7** Mean number of new variants brought by each new distinct individual added to all possible initial sample size combinations (**-A**); Number of high-quality variant positions per 100 base pair (bp) across 852 gene fragments ranked by their length (bp), overall and for each species (**-B**).

**Fig. S8** Comparison of nucleotide diversity (*theta.pi*) distributions between main cpDNA lineages (B and A or C) for *Q. robur* (586 genes) and *Q. petraea* (449 genes). The histogram represents lineage B for *Q. robur*. Data are available in both lineages within each species for at least 8 gametes per lineage, and a minimum of 200 bp per gene fragment.

**Table S1** Description of amplicons: primer sequences, original candidate gene list, targeted biological functions (see references), candidate gene type, fragment expected size and position in the *orict* original working assembly, preliminary results based nucleotide quality for obtained sequences, and validation decision after excluding paralog amplification.

**Table S2** Functional annotation results from Blast2GO (-A), comparison of BlastX best hits results (according to *E-values*) between consensus sequences of the *orict* working assembly and the *ocv4* assembly (-B), and comparisons of BlastN results of consensus sequence for both *orict* and corresponding expected amplicon (*orict-cut)* onto *ocv4* (-C).

**Table S3** Description of all variants single base positions, with sample sizes, alleles, genotypes counts, various statistics, and generic format for genotyping essays input data. Species samples exclude the 2 most introgressed individuals.

**Table S4** Description of all polymorphisms as in Table S3, but with a characterization of the length, sequence motifs, contiguous base positions for complex polymorphic regions including indels, SNPs and SSRs (see also Table S5 for SSR positions).

**Table S5** SSR patterns as detected from the *mreps* software.

**Appendix S1** Additional method details.

**Appendix S2** Contigs of the original working assembly used for selecting candidate gene regions and design amplicon primers, including consensus sequences and reads where nucleotides with Phred score below 20 have been masked.

**Appendix S3** Sequences of chosen contig consensus and singletons sequences for functional annotation analyses.

**Appendix S4** Consensus sequences of 852 genomic regions obtained in this study for *Quercus petraea* and *Q. Robur* individuals. “(N)^9^” : represents a low-quality fragment of a length below ~1 kb separating Forward and Reverse amplicons; “n” represents positions with a majority of nucleotides with phd score below 30. “(-)^x^”: means that the insertion is a minor allele at that position, x being the size of the indel.

**Appendix S5** Nucleotide sequence data of 394 gene regions for one *Quercus ilex* individual, heterozygote sites being indicated by IUPAC codes.

**Appendix S6** Outputs from Blast2GO analyses.

## Notes

https://github.com/garniergere/Reference.Db.SNPs.Quercus

### Competing Interest Statement

The authors have declared no competing interest.

### Summary of Updates

Version 4 of this preprint has been peer-reviewed and recommended by Peer Community In Forest and Wood Sciences (https://doi.org/10.24072/pci.forestwoodsci.100003)

