## Supporting-information for "High-quality SNPs from genic regions highlight introgression patterns among European white oaks (*Quercus petraea* and *Q. robur*)": Appendix-S1.pdf

**Lang et al. 2021**

[§]Corresponding author

Pauline Garnier-Géré

INRA, UMR 1202 Biodiversity Genes & Communities, F- 33610 Cestas, France; Univ. Bordeaux, UMR 1202 Biodiversity Genes & Communities, Bordeaux, F-33400 Talence, France

### Appendix S1 - Supporting Methods

*Original assembly description (see also Figure S2, supporting information)*

Sequences from the 14 cDNA libraries were obtained from various tissues and developmental stages (bud, leaf, root and wood-forming tissues) from a total of 146 individuals identified as belonging to both species, and that were sampled in 3 different French regions (South-West, North-East and North-West). These sequences were thus likely to target a large range of expressed genes. We performed the first working assembly for those sequences, with the main aim of avoiding paralog assembly while limiting split contigs with overlapping homolog sequences (Figure S2, supporting information), we initially pre-processed all sequences by removing low-quality EST, keeping those with a PHRED score above 20 (PHRED software, Ewing *et al*. 1998) for at least 90% of base pairs (bp) within a minimum of 100 bp. Vector-related sequences were trimmed or masked using Cross_match (www.phrap.org/phredphrapconsed.html) and BLAST analyses (Altschul *et al*. 1990, 1997) against the UniVec database (https://www/ncbi.nlm.nih.gov/tools/vecscreen/univec/). The ~90 000 sequences so obtained were assembled with the STACK_PACK pipeline (Miller *et al*. 1999) with the aim of avoiding the assembly of paralogs while at the same time limiting split contigs belonging to homolog sequences. The 3 main steps followed were 1) the "loose" clustering with the d2_cluster program (Burke *et al.* 1999), 2) the contig assembly within clusters with Phrap (www.phrap.org/phredphrapconsed.html) and 3) the final alignment and consensus sequence generation using STACK_Analysis and CRAW that accounts for alternative splicing variation (Burke *et al*. 1998). An iterative PHRAP step was also used for the largest contigs (including one or two orders of magnitude more reads than the average contig size), splitting contigs when quality of alignments was poor in low-depth regions to avoid possible paralog assembly. The final assembly includes 13477 contigs and 74 singletons is given in Appendix S2 with nucleotides having a Phred scored below 20 being masked with "?" using the SeqQual pipeline (https://github.com/derfR/SeqQual/). The libraries used in this assembly have since been named A, B, F to O, and S, and were included in larger transcriptome resources for *Quercus* species (Ueno *et al*. 2010).

*Choice of fragments for re-sequencing*

Expressional and functional candidate genes information was compiled for targeting those potentially involved in white oaks' divergence and/or local adaptation (Fig. S2-B and Table S1, Supporting information). Briefly, model species databases were searched for gene accessions by gene ontology (GO) and metabolic pathways keywords. Those sequences were first Blasted (Altschul et al. 1990, 1997) against our working oak assembly. Second, the sequences from their best hits were extracted (see filtering criteria in Fig. S2-B, Supporting information) and re-Blasted against the non-redundant protein (NR) database at NCBI. Third, their annotation was compared to those of the initial gene accessions, allowing 95% of hits from the oak assembly to be validated (step 2 in Fig. S2-B, Supporting information). Expressional candidate genes sequences from bud tissues or stress treatment libraries and a random set of ESTs were also directly sampled across the oak assembly generated above (see Table S1, column F, Supporting information). Primers were designed with the OSP software (Hillier and Green 1991) by setting up homogenous melting temperatures constraints and excluding low-complexity propositions. We also checked that they were located preferentially in the 3' ends of large contigs, but in regions where putative variants were absent compared to contiguous regions that could include variants (see Step 3 in Fig. S2-B, Supporting information, and primers provided in columns V and W of Table S1, Supporting information). At this stage, we also wanted to avoid targeting more conserved genes *a priori*. Thus we visually examined, among pre-selected contigs from the oak assembly, the ~100 providing BlastX results with lowest E-values (below $<10^{-80}$), in order to compare their putative polymorphisms patterns with another set of contigs with higher E-values (~ $10^{-30}$, see Fig. S2-B, supporting information). We verified that the lowest E-values contigs did not correspond to those with the lowest numbers of polymorphisms, or with an absence of putative polymorphisms. Predicted amplicons were Blasted against each other and onto our assembly to exclude those with potential amplification problems and multiband patterns. They were also checked for their depth and presence of putative polymorphisms in contigs alignment, yielding finally 2000 amplicons for resequencing (Fig. S2-B, Supporting information).

*Preliminary analyses of the 1968 amplicons (Fig. 1-A)*

Overall, more than 85% of the designed amplicons were successful in both individuals, one from each species. We tested whether fragments amplified only in one individual or the other (called fragments A) were more polymorphic overall than those amplifying in both individuals (called fragments B): using a minimum Phred score of 30 (i.e. error rate below 0.001), we extracted fragments with a minimum length of 200 bp and maximum mean proportion of missing data of 50%, as computed across overlapping windows by 50% across fragments. Among these, a similar amount of fragments A were found in both individuals (158 in *Q. robur* 11P individual versus 155 in *Q. petraea* Qs21 individual). One SNP (or heterozygote here) per 654 bp was observed on average across the ~99 kb amplified in Q. robur/11P only, compared to one SNP per 340 bp (across ~ 500 kb) using data on the same individual for fragments B. For the Q. petraea/Qs21 individual, the same statistics are respectively: one SNP per 926 bp across ~98 kb, and one per 342 bp across ~500 kb). Filtering more strongly on quality with a maximum proportion of missing data of 25% slightly increased the number of fragments which can be considered as being amplified only for one individual or the other, but the same trend remains (although with more similar values in Q. robur/11P and Q. petraea/Qs21) of around twice less heterozygote at SNPs compared to fragments B with similar quality.

Although it is difficult to conclude on the basis of one individual per species, there is no evidence that fragments A are more polymorphic than fragments B. We also need to be prudent since with the Sanger technique used here, the quality filtering may also have masked some heterozygote indels in diploid sequences (see the part *Treatment of diploid sequences…* below for the full data obtained), and thus also subsequent parts in the fragments, yielding stretches of low-quality positions due to the frame shift of the second strand, which might also harbor polymorphic sites. Indeed, 24% of fragments A had proportion of missing data above 25%, compared to 9% for fragments B, indicating an overall lesser apparent quality and thus the possibility that some polymorphisms may have been missed. However, the same treatment was used for Fragments A and B here, and thus polymorphic sites may have been missed also in fragments B. Overall, we can consider that given the strategy followed for choosing the fragments and designing the primers, given the preliminary results above on the 1968 amplicons, and given the results showing a large nucleotide diversity overall in these species (see Results of the main text), there is no strong evidence that we targeted genic regions that were particularly conserved.

*Functional annotation of re-sequenced genic regions using BlastX and BLAST2GO analyses*

BlastX search (using BlastX 2.6.0+ program at NCBI (https://blast.ncbi.nlm.nih.gov/Blast.cgi) was finally performed on 396 sequences from *orict* (our working assembly original contigs) and 368 sequences from the most recent oak assembly *ocv4* (Lesur *et al*. 2015, and see Table S2-A,-B,-C and Appendix S3, Supporting information, for contig consensus sequences), based on their annotation consistency, BlastN and BlastX lowest *E-values* and highest % of identity, consensus length (below 6 kb) and a minimum number of IUPAC ambiguity codes (below 50) scattered across sequences. We excluded ~60 *ocv4* consensus sequences (~8%) with stretches of such codes (from around 50 to 430 bp) that indicate possible paralogs or alternative spliced exons in the *ocv4* assembly (Lesur *et al*. 2015, and see column P "*nb.pol.2c"* in Table S2-B, Supporting information).

For Blast2GO analyses, based on the studied genes hits similarity distribution (Appendix S6-B, Supporting information), we used 2 cut-off values: 55% for similarity and 33 (~100 bp) for high scoring segment pair (HSP). This allowed retrieving Genbank identifiers and corresponding Gene Ontology (GO) terms, which were mainly from the UniProtKB and TAIR databases. In order to examine the relevance of the original gene lists in targeting broad functional traits (column F in Table S1, Supporting information), we tested whether they contained an enrichment of particular GO terms in comparison to the randomly chosen contigs, using GO data across all sequences in each gene list.

*Treatment of diploid sequences obtained in the discovery panel with SeqQual for polymorphism discovery*

Sequence data of amplicons from the same original contig were assembled together and consensus sequences were obtained with the Phred/Phrap (www.phrap.org) suite of programs called by SeqQual. Ambiguous codes not detected as valid polymorphisms by Polyphred (https://droogs.gs.washington.edu/polyphred/) were considered as missing data and masked. Overall, the time needed to call polymorphisms with SeqQual scripts and to validate by visual examination the traces or alignments in amplicons identified with possible problems was much smaller (by a factor of at least 50) than the time needed to correct data in BIOEDIT (Hall 1999) or CodonCode Aligner (CodonCode Corporation, www.codoncode.com/aligner/ and see parameter examples at https://github.com/garniergere/SeqQual/tree/master/SeqQual_shell_ex). Using print_source-SNP-statistic.pl script output for diploid sequences (https://github.com/garniergere/SeqQual/tree/master/SeqQual_pdf/SeqQual-part3-fastools-usage.pdf), we could easily point to amplicons with simple insertion-deletion polymorphisms (*indel*s), more complex *indel* patterns that included simple sequence repeats (*SSRs*), and rare or outlying patterns such as heterozygote excess or deficit (see Results). Visual examination then allowed alignments in those complex regions to be corrected if needed. From mismatch cases that were revealed automatically when merging forward and reverse amplicons, we confirmed that more than 99% of identified heterozygotes were correct with chosen Polyphred parameter values (60 and 90 for threshold overall and genotype scores respectively), and so their absence in one of the alternative strand was due to a clear but too weak second peak for a positive heterozygote call. Also, cases of heterozygote excess that mostly showed double peaks (*DoP*) were considered as paralog amplifications and excluded (column L in Table S1, Supporting information and Fig. 1-D). Additionally for each diploid sequence with a clear automatic heterozygote indel (*HI*) pattern (i.e. a trace with mostly single peaks becoming clear DoP after a particular position and the presence of at least one homozygote individual for the deletion at that position, or with *DoP* patterns that were consistent with a particular deletion), we coded the heterozygotes at the first position with corresponding IUPAC codes (http://www.chem.qmul.ac.uk/iubmb/misc/naseq.html), followed by missing data. This allowed minimizing the amount of missing data produced by superimposed allele traces. In several cases, a second *HI* further along the sequence allowed getting back a clear open reading frame and diploid sequence information. The putative rare variants in some individuals that we missed would be located in those DNA stretches assigned to missing data due to overlapping traces after such *HI* recoded positions. These *HI* positions, because there were coded and characterized in the lists provided, homopolymers excepted for thus allow more accurate diversity estimates (Tables S3 and S4, supporting information).

*Treatment of homopolymers in diploid sequences*

Homopolymers, mostly from 8 to 10 A or T repeats depending on the amplicons, often stopped the correct functioning of the polymerase during sequencing after their positions and were observed in 104 amplicons (Tables S1 and S5, supporting information, for their description, position and presence across genes), so heterozygote indels in those regions were generally masked before polymorphism counts.

*STRUCTURE analyses and runs*

We used STRUCTURE v2.3.3 (Pritchard *et al*. 2000, Falush *et al.* 2003) to infer genetic clusters and test for possible levels of introgression across individuals. Following recommended defaults, we used the admixture model allowing for mixed ancestry and the correlated allele frequencies assumption for closely related populations. We first drew one polymorphic locus at random per genic region and simulated 10 replicates for each value of K clusters (1 to 5) with burn-in and post- burn-in periods of 100,000 and 1,000,000 iterations respectively. Due to very low standard deviation across replicates of the data log likelihood given K (ln Pr(X/K)), we further tested the robustness of the results to genetic stochasticity by resampling loci at random for each of 10 replicate datasets in 3 different manners: 1) one per region, 2) one per 100 bp block, and 3) one per 200 bp block along genes. The block sizes were chosen to sample more loci while keeping low levels of background linkage disequilibrium (LD) *a priori*, given the already high number of independent gene regions (>800), and given that STRUCTURE permits the inclusion of weakly linked markers (Falush *et al*. 2003). Examples of STRUCTURE data and parameter files are archived as recommended by Gilbert *et al.* (2012) along with R scripts for plots including Bayesian confidence intervals in the K=2 case **(**https://github.com/garniergere/Reference.Db.SNPs.Quercus/tree/master/STRUCTURE.files**).** Missing data were below 20% across loci, at least 12 gametes were present in the original morphological species, and an arbitrary *maf* of at least 9% across polymorphisms allowed singletons to be excluded. We examined both ln(Pr(X/K)) and ΔK (Evanno *et al*. 2005) statistics using STRUCTURE HARVESTER (Earl and von Holdt 2012).
