## Supporting-information for "High-quality SNPs from genic regions highlight introgression patterns among European white oaks (*Quercus petraea* and *Q. robur*)": Appendix-S6-A-B.docx

**Appendix S6** Outputs from Blast2GO analyses: A) Species distribution across BlastX hit results for the original contig consensus sequences list (see Appendix S3 and BlastX parameters in Table S2). B) Distribution of sequence similarity percentages across BlastX hit results for the same contig consensus sequences list (Appendix S3).

**A)**

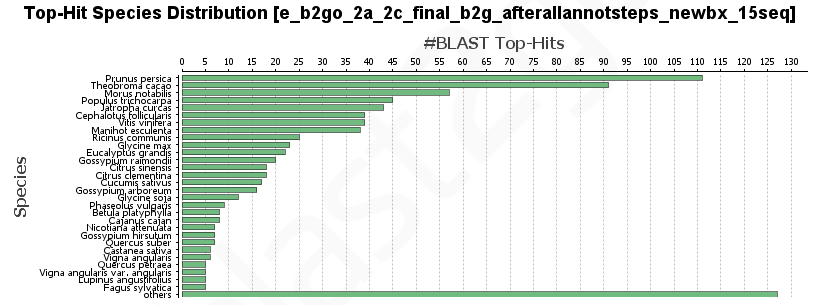

**B)**

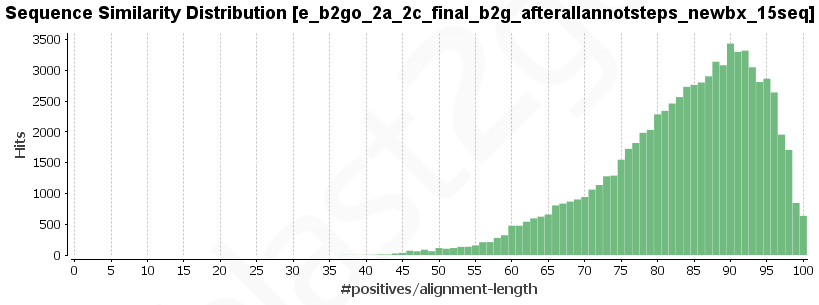
