## Supporting-information for "High-quality SNPs from genic regions highlight introgression patterns among European white oaks (*Quercus petraea* and *Q. robur*)": Supporting-information-Fig-S1-to-S8.pdf

Suporting information for **Lang et al.**

Figures S1 to S7

**Fig. S1** Sampling site locations within the natural geographic distribution of *Q. petraea* and *Q. robur*. Vector map is from *http://www.naturalearthdata.com* and distribution areas from Euforgen (*http://www.euforgen.org/distribution-maps/*)

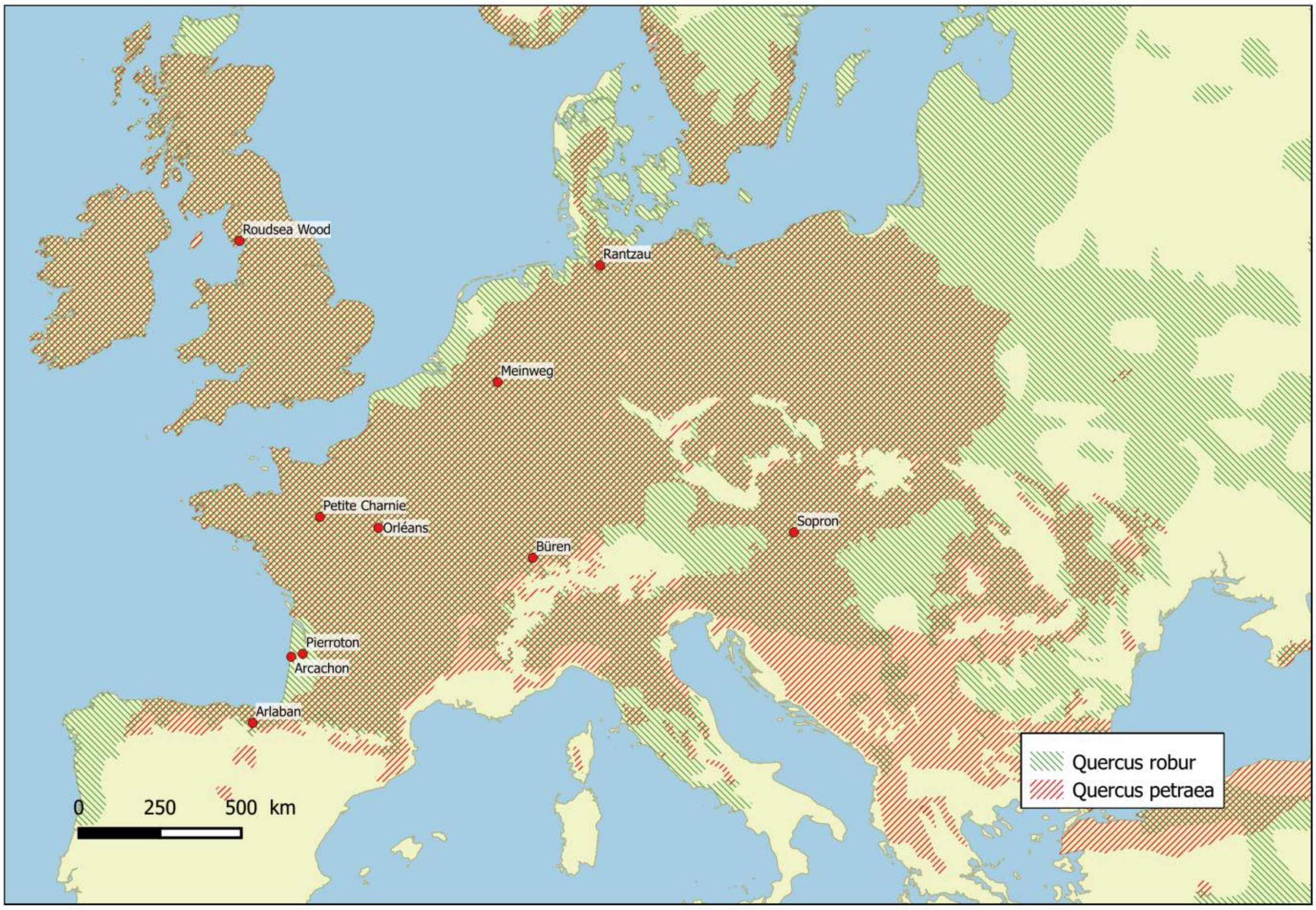

**Fig. S2** Working assembly steps and softwares (A), and bioinformatic strategy for search of candidate genes and amplicon choice (B).

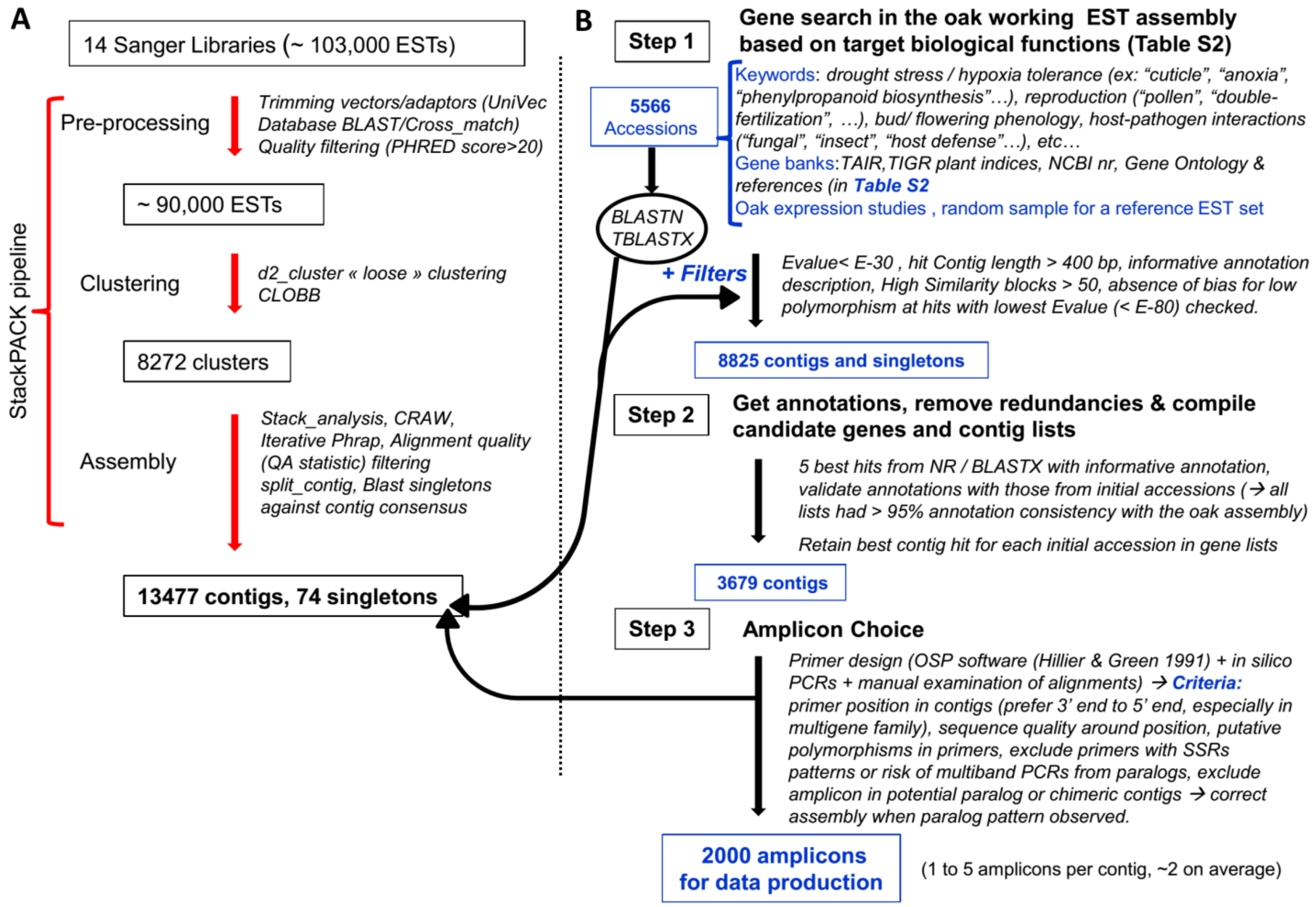

**Fig. S3** Plots of the ΔK values from the Evanno *et al*. (2005) method (S3-A, -B, -C, -D, -E), and of the mean values of the estimated probability ln (of the data given K) with standard deviations for K ranging from 1 to 5 (S3-F to S3-J), which show support for K=2. Plots are from the STRUCTURE HARVESTER program.

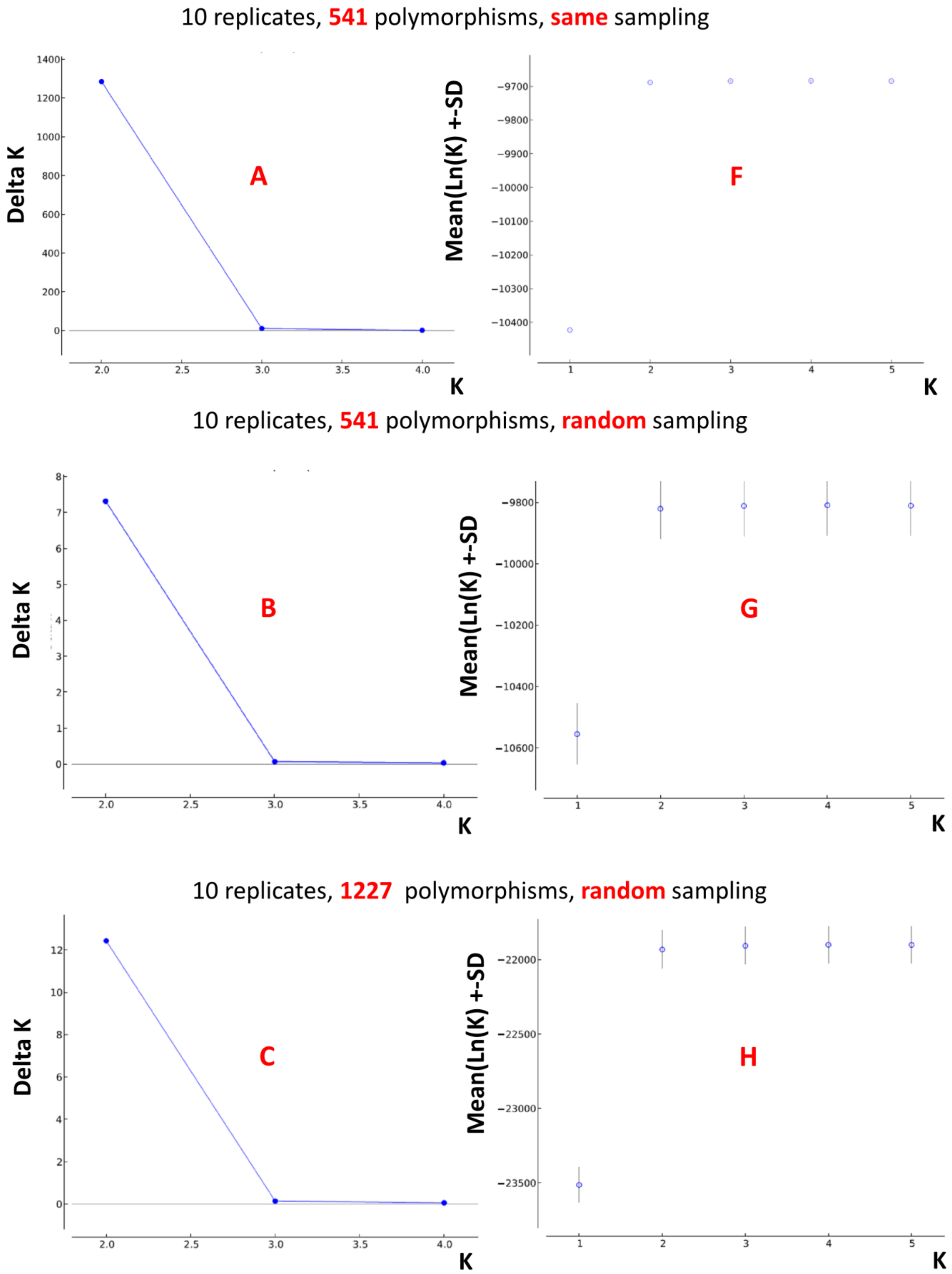

10 replicates, **1785** polymorphisms, **same** sampling

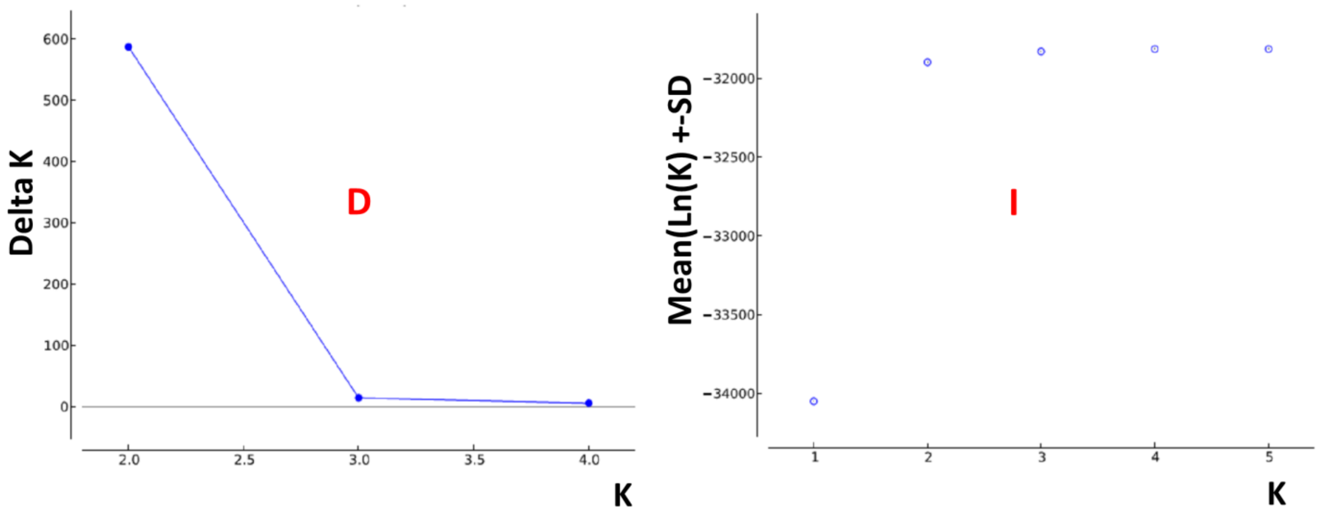


10 replicates, **1785** polymorphisms, **random** sampling

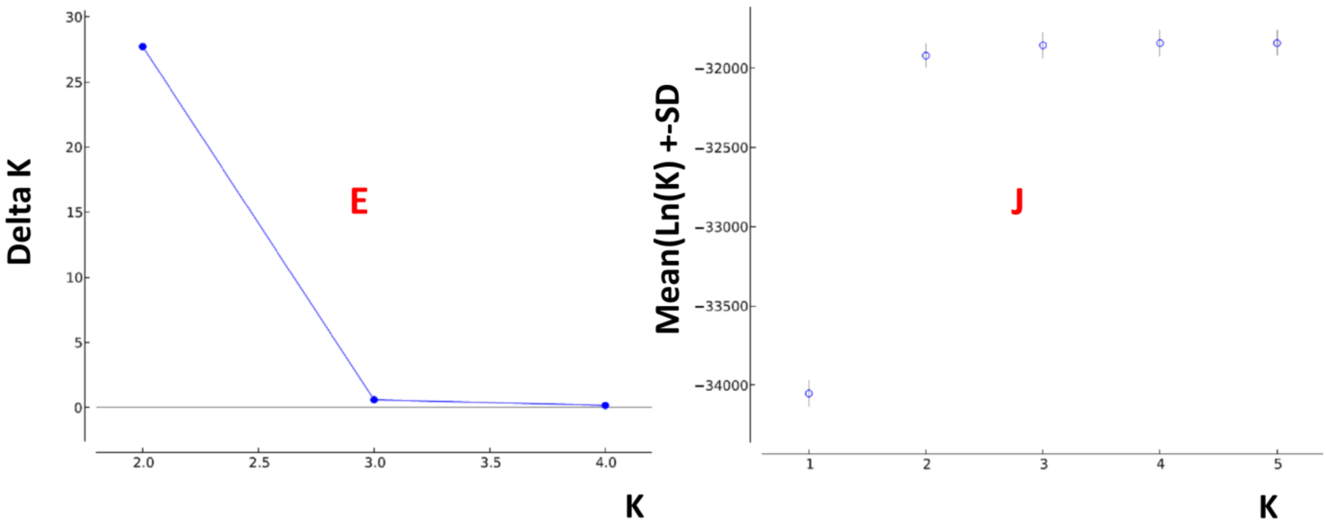

**Fig. S4** Distributions of Gene Ontology (GO) terms for the consensus sequences in Appendix S3, at level 2 (-A, -B, -C) and level 3 (-D, -E, -F): A- and D- for Biological Process, B- and E- for Molecular Function, C- and F- for Cellular Component. Annotation rules: E-value<$10^{-30}$, annotation cut-off 70, GO weight 5, HSP coverage cutoff 33%. Filtering applies for at least 5 sequences and a node score of 5 per GO term (but see rare exceptions in Table S2).

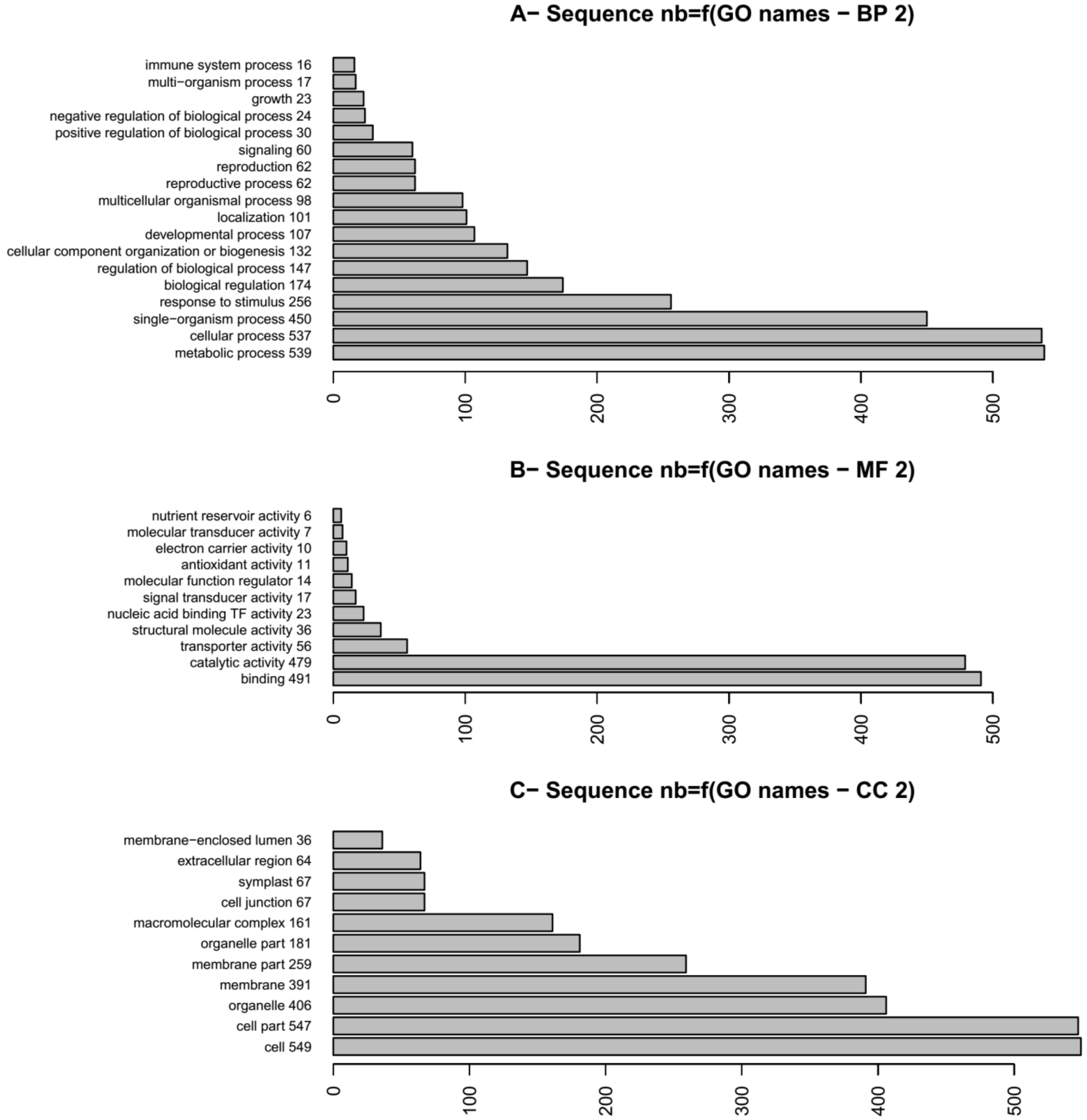

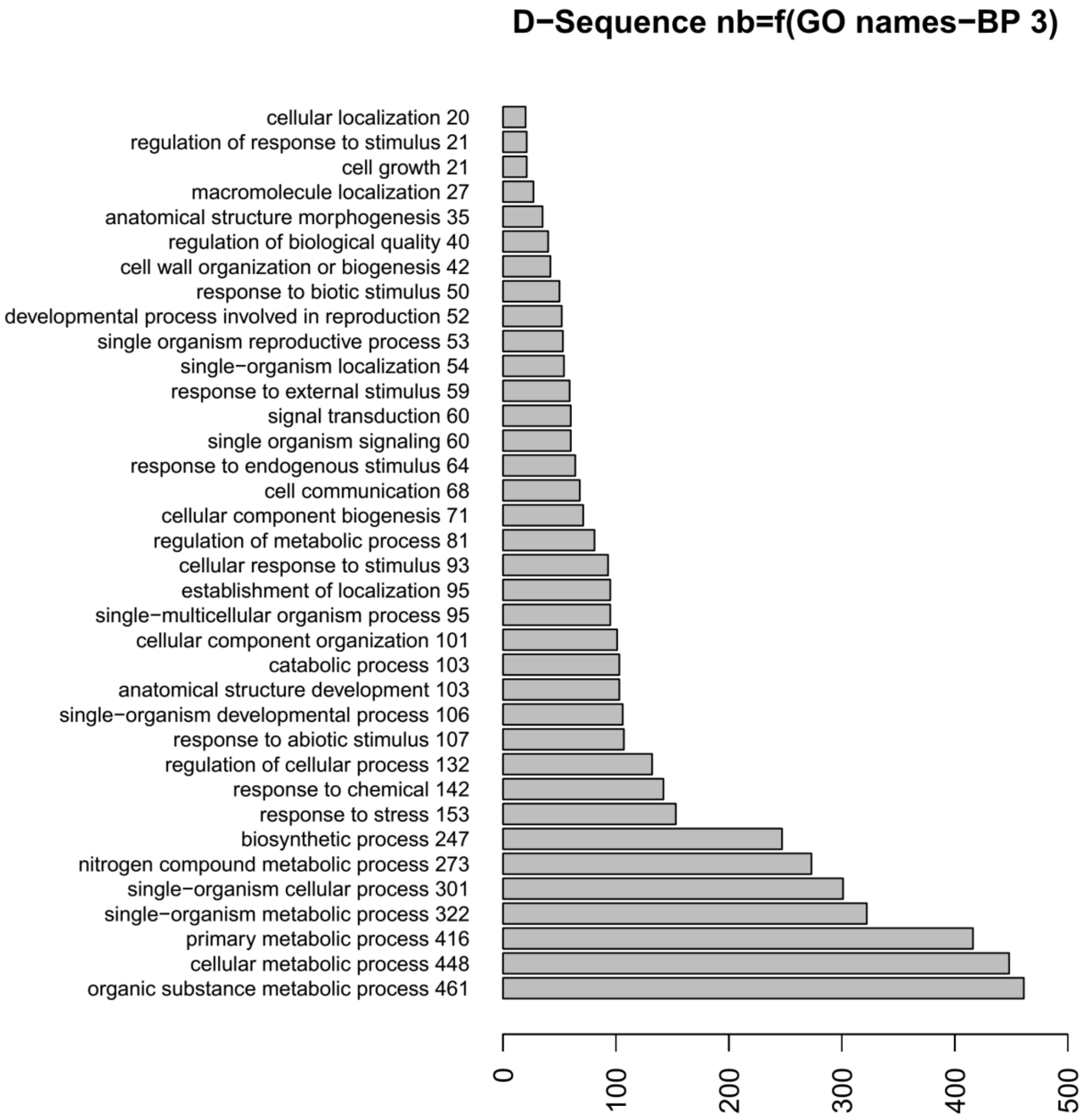

D−Sequence nb=f(GO names−BP 3)
cellular localization 20
regulation of response to stimulus 21
cell growth 21
macromolecule localization 27
anatomical structure morphogenesis 35
regulation of biological quality 40
cell wall organization or biogenesis 42
response to biotic stimulus 50
developmental process involved in reproduction 52
single organism reproductive process 53
single−organism localization 54
response to external stimulus 59
signal transduction 60
single organism signaling 60
response to endogenous stimulus 64
cell communication 68
cellular component biogenesis 71
regulation of metabolic process 81
cellular response to stimulus 93
establishment of localization 95
single−multicellular organism process 95
cellular component organization 101
catabolic process 103
anatomical structure development 103
single−organism developmental process 106
response to abiotic stimulus 107
regulation of cellular process 132
response to chemical 142
response to stress 153
biosynthetic process 247
nitrogen compound metabolic process 273
single−organism cellular process 301
single−organism metabolic process 322
primary metabolic process 416
cellular metabolic process 448
organic substance metabolic process 461
0
100
200
300
400
500

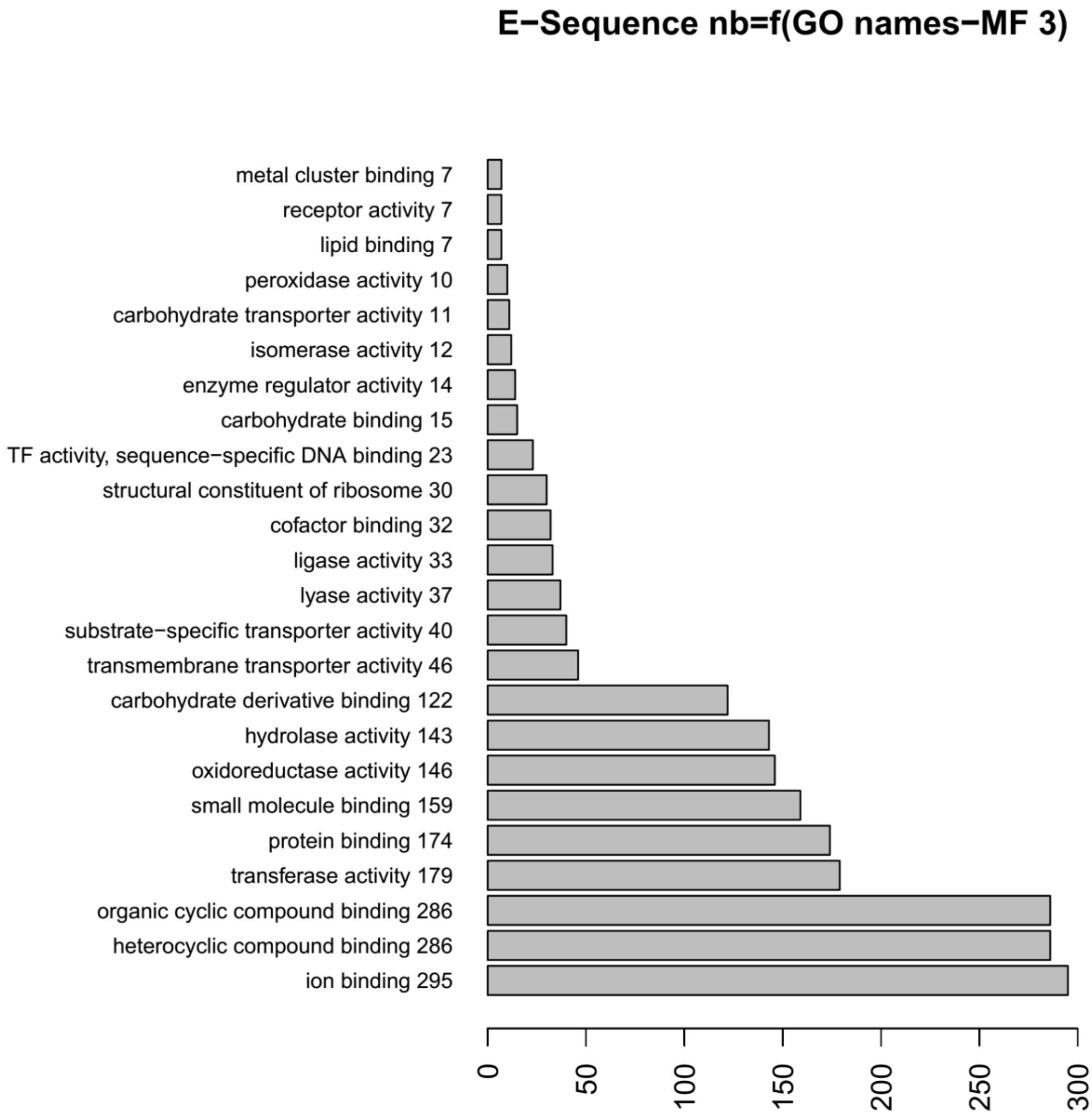
E−Sequence nb=f(GO names−MF 3)
metal cluster binding 7
receptor activity 7
lipid binding 7
peroxidase activity 10
carbohydrate transporter activity 11
isomerase activity 12
enzyme regulator activity 14
carbohydrate binding 15
TF activity, sequence−specific DNA binding 23
structural constituent of ribosome 30
cofactor binding 32
ligase activity 33
lyase activity 37
substrate−specific transporter activity 40
transmembrane transporter activity 46
carbohydrate derivative binding 122
hydrolase activity 143
oxidoreductase activity 146
small molecule binding 159
protein binding 174
transferase activity 179
organic cyclic compound binding 286
heterocyclic compound binding 286
ion binding 295
0
50
100
150
200
250
300

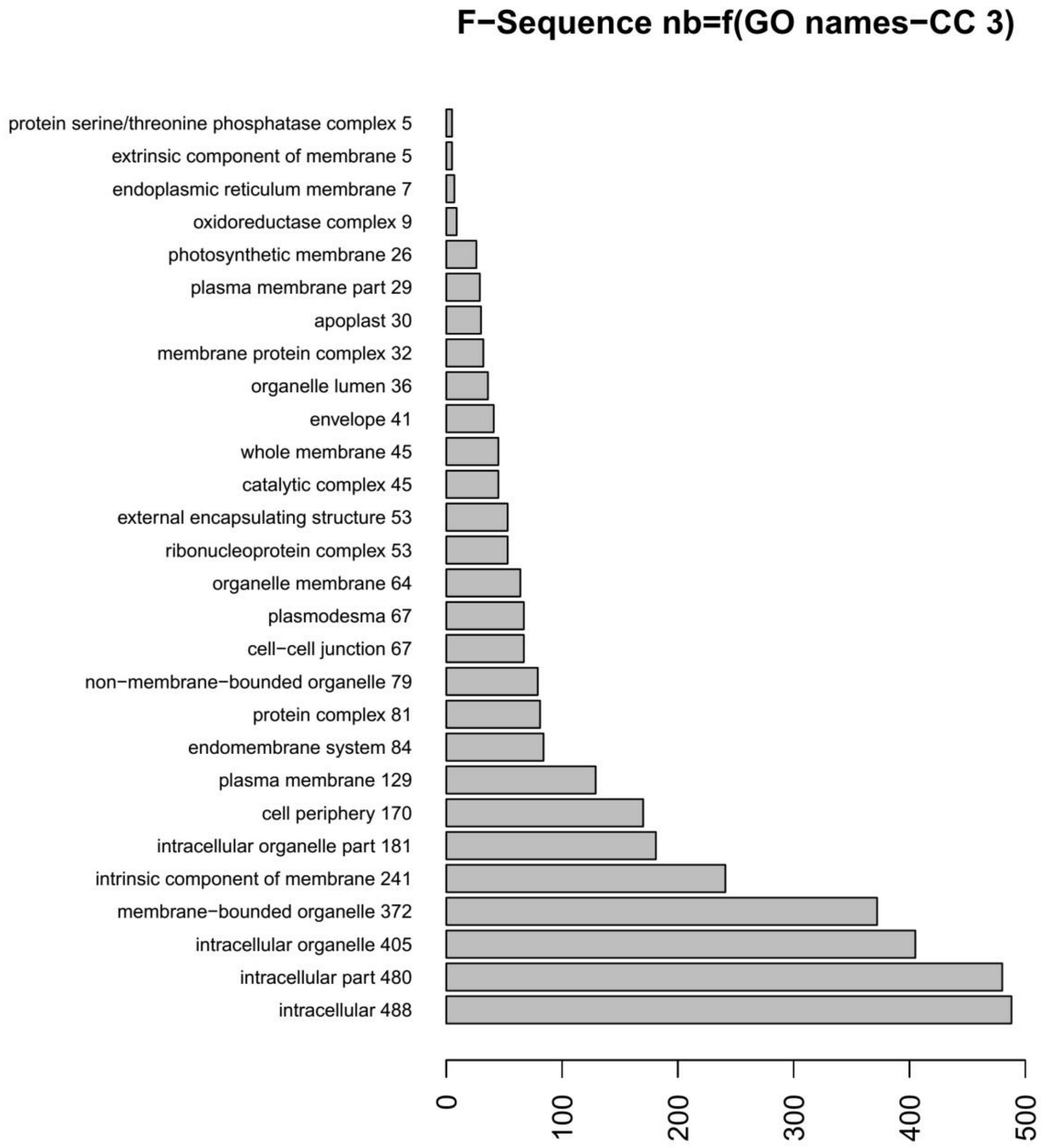
F−Sequence nb=f(GO names−CC 3)
protein serine/threonine phosphatase complex 5
extrinsic component of membrane 5
endoplasmic reticulum membrane 7
oxidoreductase complex 9
photosynthetic membrane 26
plasma membrane part 29
apoplast 30
membrane protein complex 32
organelle lumen 36
envelope 41
whole membrane 45
catalytic complex 45
external encapsulating structure 53
ribonucleoprotein complex 53
organelle membrane 64
plasmodesma 67
cell−cell junction 67
non−membrane−bounded organelle 79
protein complex 81
endomembrane system 84
plasma membrane 129
cell periphery 170
intracellular organelle part 181
intrinsic component of membrane 241
membrane−bounded organelle 372
intracellular organelle 405
intracellular part 480
intracellular 488
0
100
200
300
400
500

**Fig. S5** Distributions of GO terms across different gene lists (*bud*, *abiotic* and *biotic*) at Biological Level 2, and Fisher exact tests across pairs of sequence clusters with the same GO terms between the random list and other lists. Significance levels *: P<0.05.

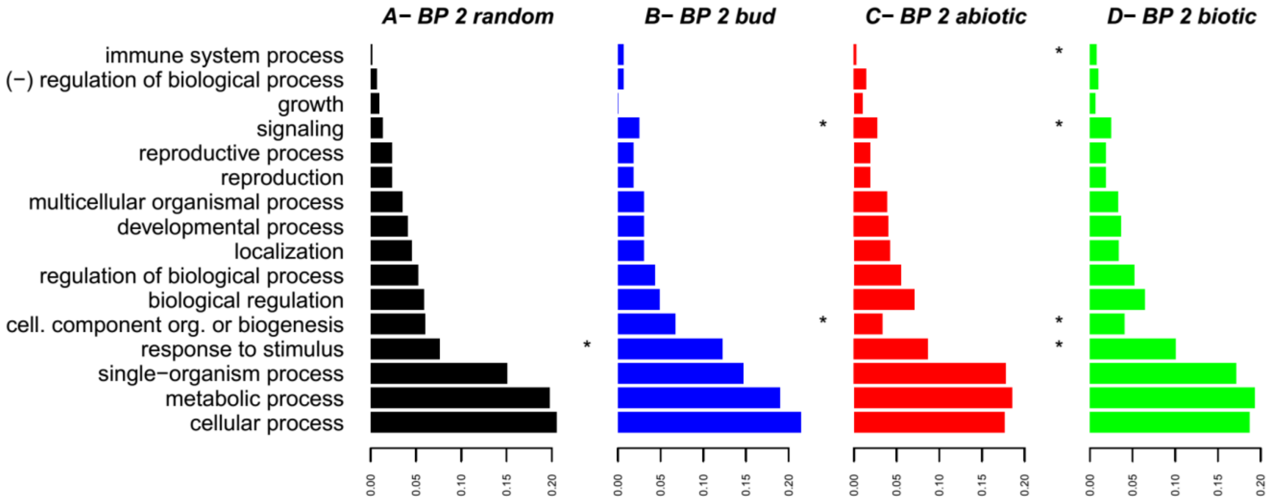


**Fig. S6 A-J** Posterior assignment probabilities (*Q*-values) of 24 individuals attributed to 2 clusters (STRUCTURE analysis) for different numbers of polymorphisms, different sampling of SNP data, and different plots of credible intervals.

*A to D: 541 polymorphisms across 541 gene regions with either the same variant positions (A and B) or a random sample of variants for 10 replicates (C and D). E to H: 1785 polymorphisms across 541 gene regions split into 100 bp blocks, with either the same variants (E and F) or a random sample of variants across 10 replicates (G and H). I and J: 1227 polymorphisms across 541 gene regions split into 200 bp blocks with a random sample of variant positions across 10 replicates. Probabilities are sorted in increasing order of belonging to one cluster or the other (Q. robur is indicated by "qr" across individuals' names and Q. petraea by "qp". Each bar is one sampled individual with its mean Q-value and associated 90% Bayesian confidence intervals (BCI). BCI are computed either with the mean upper and mean lower bound values across replicates (A, C, E, G and I) or as maximum and minimum bound values across replicates (B, D, F, H and J).*

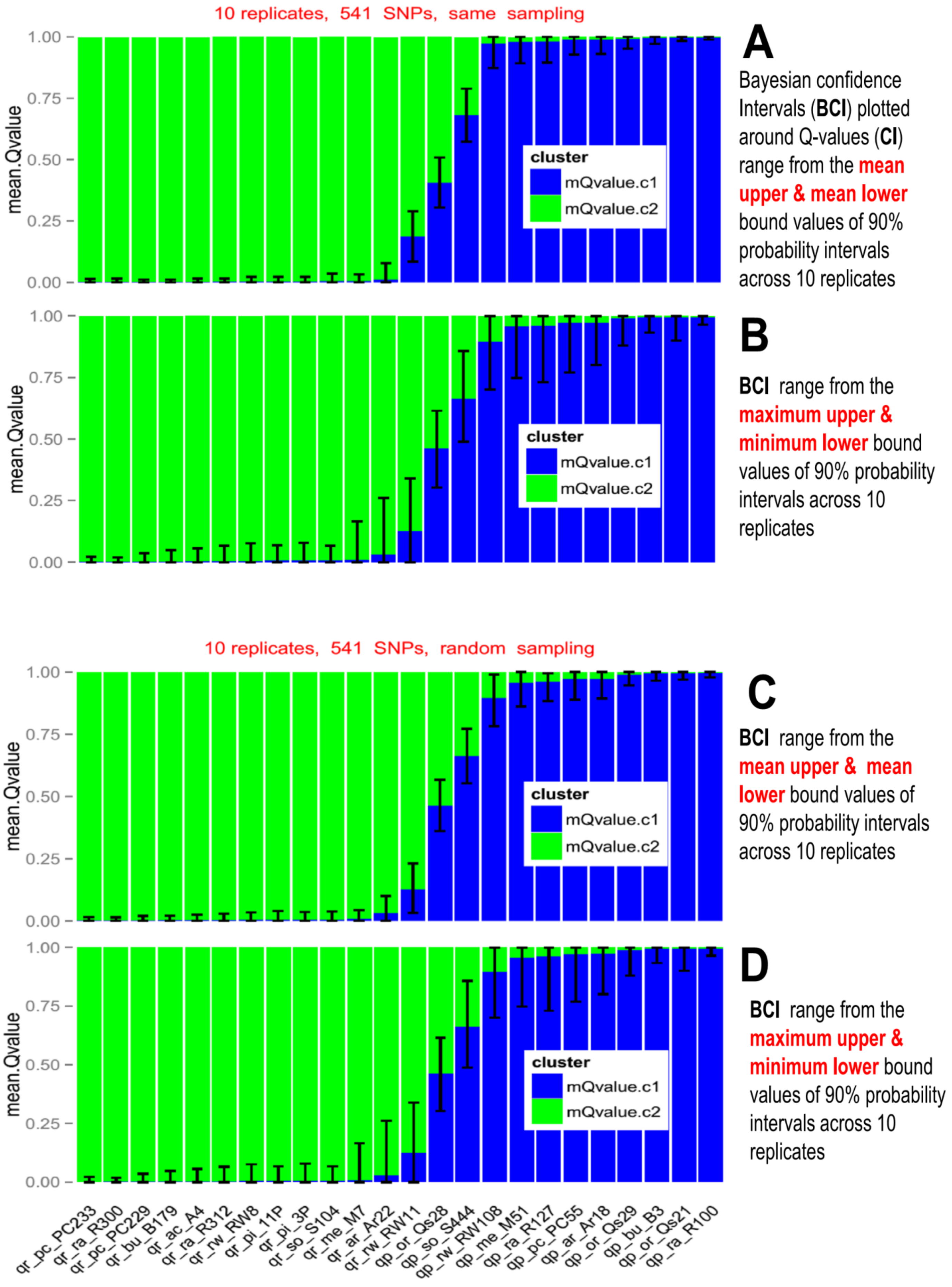
10 replicates, 541 SNPs, same sampling
A
Bayesian confidence Intervals (BCI) plotted around Q-values (CI) range from the mean upper & mean lower bound values of 90% probability intervals across 10 replicates
B
BCI range from the maximum upper & minimum lower bound values of 90% probability intervals across 10 replicates
10 replicates, 541 SNPs, random sampling
C
BCI range from the mean upper & mean lower bound values of 90% probability intervals across 10 replicates
D
BCI range from the maximum upper & minimum lower bound values of 90% probability intervals across 10 replicates
mean.Qvalue
1.00
0.75
0.50
0.25
0.00
cluster
mQvalue.c1
mQvalue.c2
qr_pc_PC233
qr_ra_R300
qr_pc_PC229
qr_bu_B179
qr_ac_A4
qr_ra_R312
qr_rw_RW8
qr_pi_11P
qr_pi_3P
qr_so_S104
qr_me_M7
qr_ar_Ar22
qr_rw_RW11
qp_or_Qs28
qp_so_S444
qp_rw_RW108
qp_me_M51
qp_ra_R127
qp_pc_PC55
qp_ar_Ar18
qp_or_Qs29
qp_bu_B3
qp_or_Qs21
qp_ra_R100

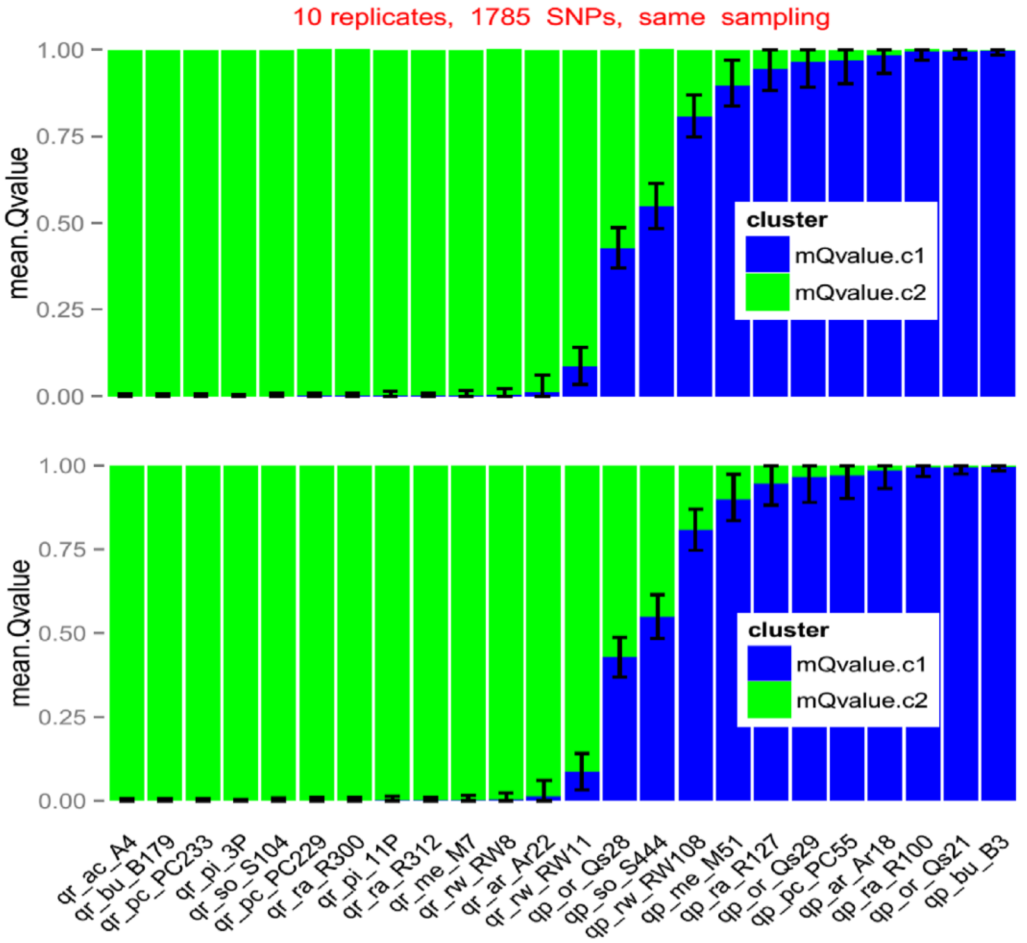


**E**

**BCI** range from the **mean upper & mean lower** bound values of 90% probability intervals across 10 replicates

**F**

**BCI** range from the **maximum upper & minimum lower** bound values of 90% probability intervals across 10 replicates

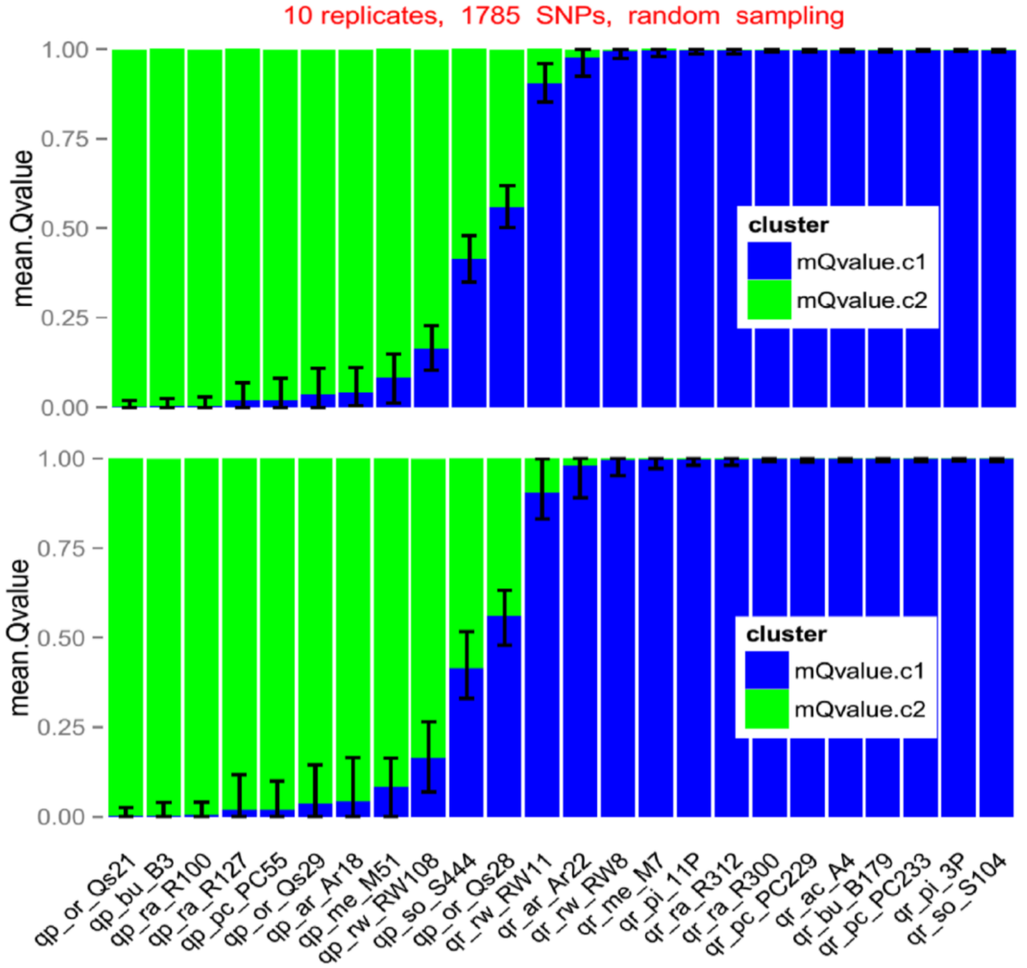


**G (=Fig.4)**

**BCI** range from the **mean upper & mean lower** bound values of 90% probability intervals across 10 replicates

**H**

**BCI** range from the **maximum upper & minimum lower** bound values of 90% probability intervals across 10 replicates

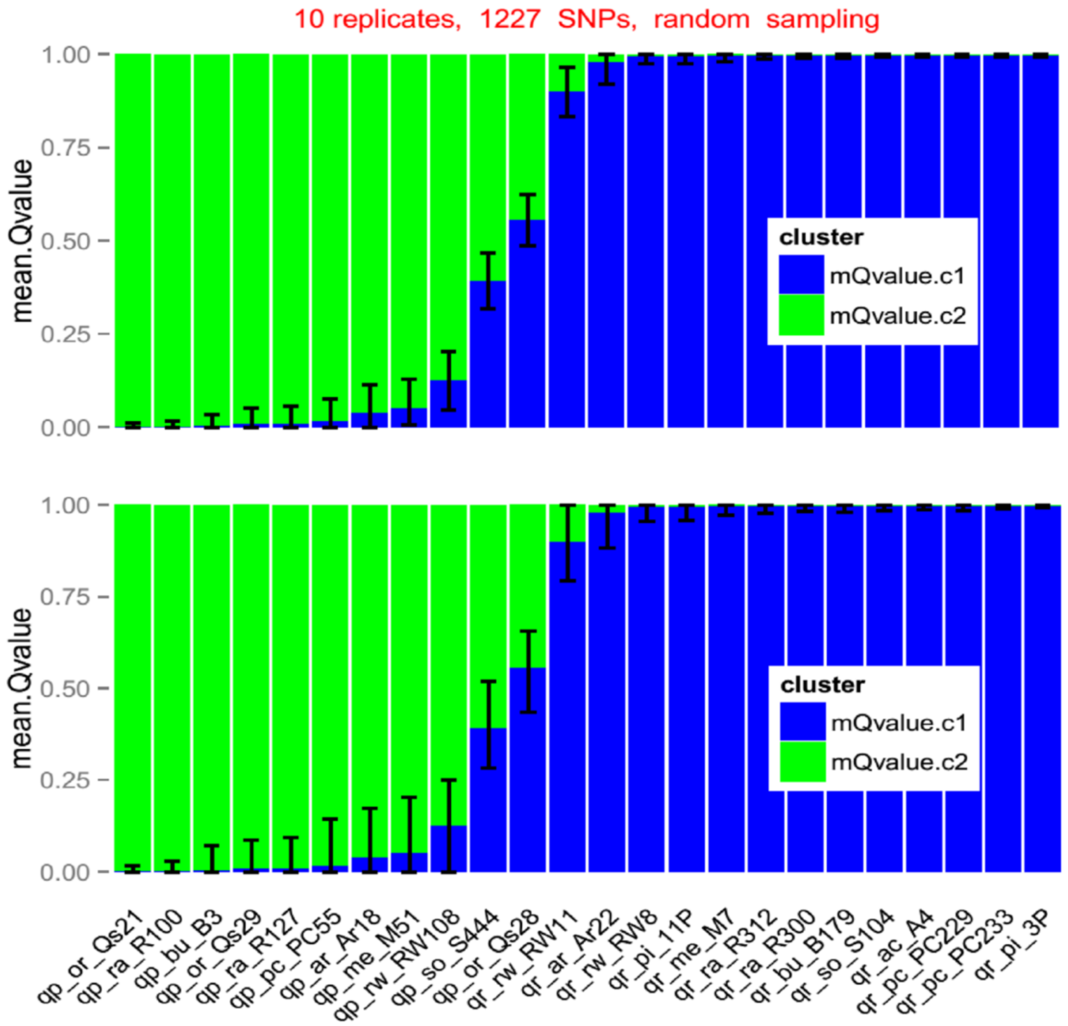


**I**

**BCI** range from the **mean upper & mean lower** bound values of 90% probability intervals across 10 replicates

**J**

**BCI** range from the **maximum upper & minimum lower** bound values of 90% probability intervals across 10 replicates

**Fig. S7** Mean number of new variants brought by each new distinct individual added to all possible initial sample size combinations (-A); Number of high-quality variant positions per 100 base pair (bp) across 852 gene fragments ranked by their length (bp), overall and for each species (-B).

Figure S7-A

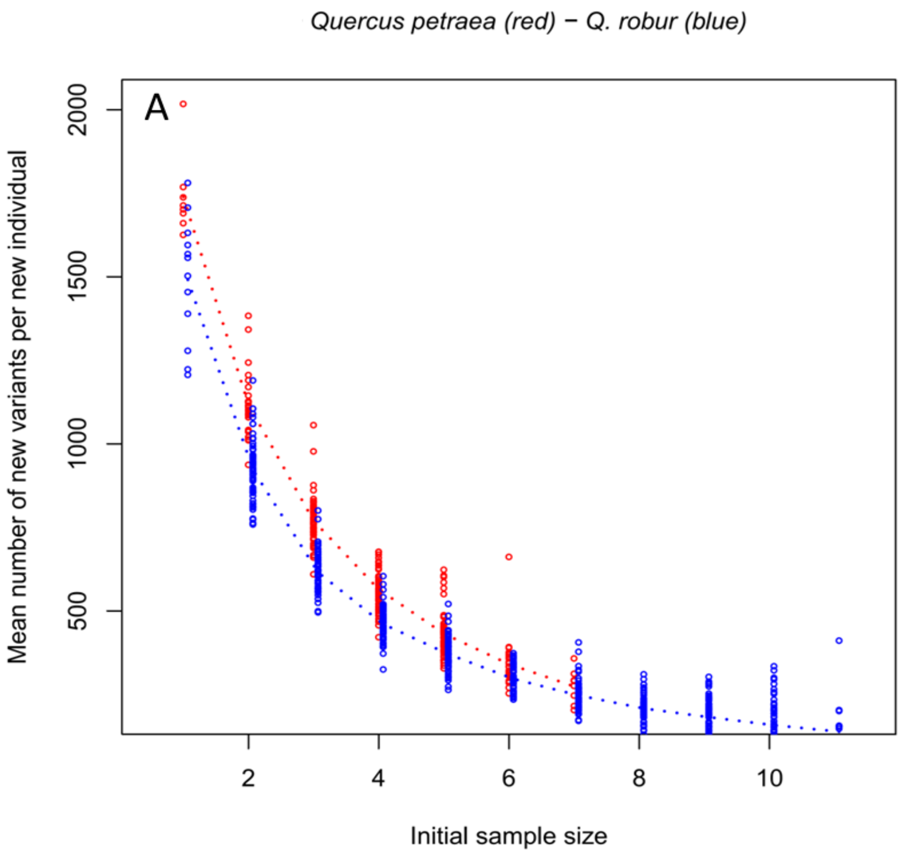


Figure S7-B

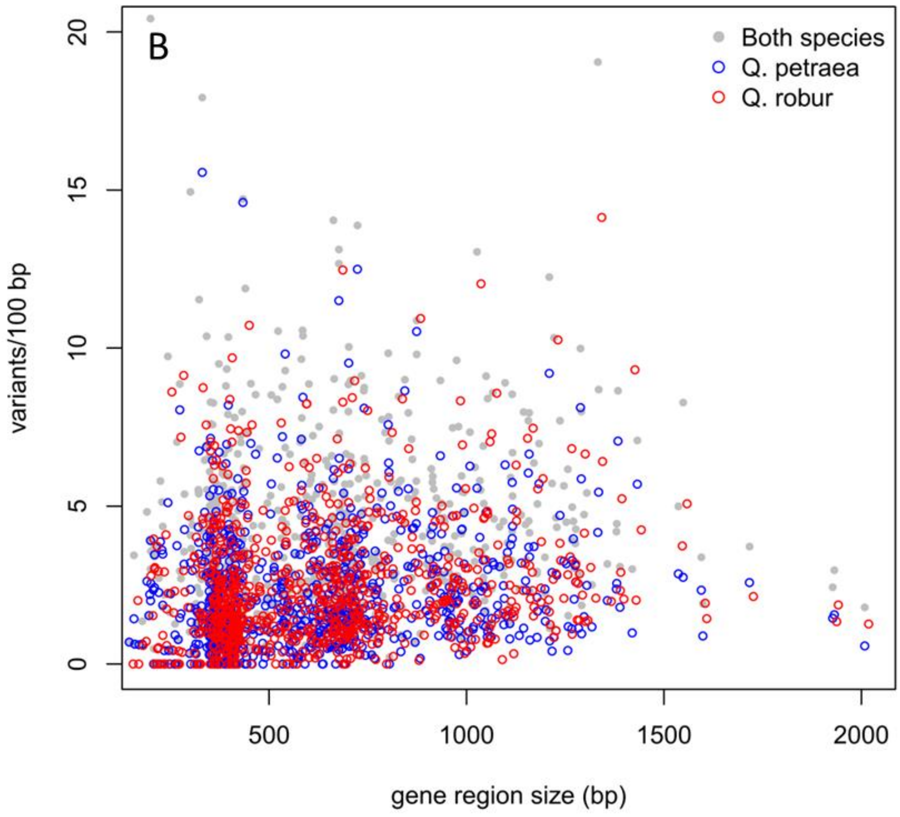

Fig. S8

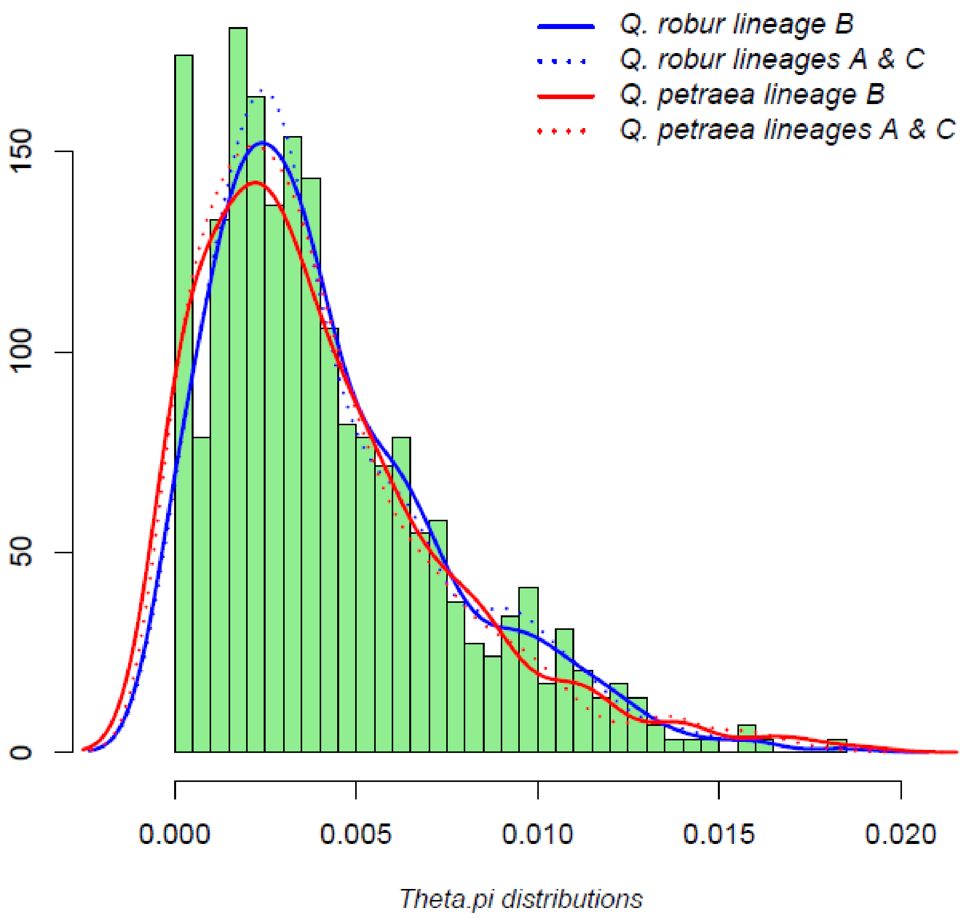
Q. robur lineage B
Q. robur lineages A & C
Q. petraea lineage B
Q. petraea lineages A & C
0
50
100
150
0.000
0.005
0.010
0.015
0.020
Theta.pi distributions
